# The neural mechanisms of concept formation over time in music

**DOI:** 10.1101/2024.11.06.622228

**Authors:** L. Bonetti, G. Fernández-Rubio, S.A. Szabó, F. Carlomagno, P. Vuust, M.L. Kringelbach, E. Brattico

## Abstract

The interplay between abstraction, learning and memory is crucial for humans to interact with the environment. Abstraction in relation to concept formation is typically studied with Gestalt stimuli varying in their physical properties while maintaining abstract internal relations. The role of temporal integration in recognising abstract concepts has been, however, overlooked even in a sensory domain relying on temporal item succession such as audition. Here, we investigated the neural mechanisms underlying abstract concept generalisation over time using a musical adaptation of the old/new recognition paradigm. During magnetoencephalography (MEG) recordings, 108 participants categorised five-tone sequences as previously learned or novel or as musical or not. The sequences varied in pace, ranging from very fast (125 ms per sound) to very slow (5000 ms per sound). On the behavioural level, sequence duration exerted a substantial impact on the recognition and musicality judgments of the novel sequences. The differences in neural responses between memorised and novel sequences especially involved regions in the auditory cortex, medial cingulate gyrus and medial temporal lobe and were most pronounced at the highest musical pace, diminishing proportionally as the pace decreased. These findings contribute to our understanding of how the brain generalises and derives meaning from abstract concepts that unfold over time learned through the auditory system.

## Introduction

To optimise information processing, humans have evolved to rely on previous learning to understand novel information (Pearl, 2018; Sommer, 2016). A key strategy in this process is concept generalisation, a topic that has intrigued scholars for millennia, starting with the ancient Greek philosopher Aristotle, who formulated early theories of categorisation (Bronstein, 2016) and ending up with neuroscience and artificial intelligence (Geirhos et al., 2020, Sharaev et al., 2018, Wu et al., 2018). Currently, concept generalisation refers to recognising novel stimuli as members of a category based on past encounters and learning. This ability is fundamental to making inferences about the world. For example, in the interpretation of semantics, it involves recognising a new black cat as a cat after having previously seen an orange one. This also aligns with Gestalt laws, which provide a framework for understanding how humans organise and perceive complex stimuli, forming a new concept that is greater than the sum of its parts. These laws suggest that perception is organised into hierarchical layers, allowing stimuli to be divided into components without compromising overall recognition (Terhardt, 2016).

Among the several fields which contributed to the topic, cognitive psychology has arguably provided the major advances by exploring the visual, bottom-up, and knowledge-based, top-down dimensions of concept generalisation (Hasson et al., 2015). At its core, the key research question revolved around the organisation of information within the mind (Kiefer & Pulvermüller, 2012), defining ‘category’ as a structural unit encompassing multiple concepts with shared similarities (e.g. Freedman & Assad, 2016).

Cognitive scientists conducted studies in which participants were tasked with categorising encountered objects, encompassing both living and non-living entities. These objects possessed multiple properties, including visual, auditory, tactile, and olfactory dimensions, all contributing to the construction of meaning. Thus, it is generally acknowledged that the information utilised in this initial phase of research is multimodal, and the concepts employed are not abstract ones. In the pioneering studies of Rosch throughout the 1970s, the primary focus was on colours, forms, and everyday language usage (e.g., naming animals, furniture), with experimental paradigms involving the rating of items based on their ‘liked-ness’ (Rosch & Lloyd, 1978). These studies have been comprehensively summarised, critically discussed, and applied to the auditory domain of perception in the work of Hantschel & Bullerjahn (2016).

More recently, novel methodologies such as neuroimaging have been employed for comprehending not only the behavioural aspects but also the underlying brain mechanisms of concept generalisation and learning (Frixione & Lieto, 2012).For instance, Bowman & Zeithamova (2018) designed a task wherein participants categorised abstract visual concepts while undergoing functional magnetic resonance imaging (fMRI) scans. Focusing on two specific brain regions, the hippocampus and the ventromedial prefrontal cortex (VMPFC), their findings demonstrated that 75% of participants predominantly relied on prototype information. In essence, the preferred method for categorisation and generalisation involved abstracting the introduced concepts rather than retaining individual exemplars in memory.

As an additional proof of concept, Vaidya et al. (2021) conducted an fMRI study, with the aim of mapping brain regions involved in the generalisation of task structures. This entailed a training phase, through which participants learned to execute a decision-making task requiring the matching of contexts and categories, followed by completing the same task on unknown contexts and categories, without receiving feedback. Thus, they were required to generalise the information learned during the training phase. The neural regions identified during the task included the orbitofrontal cortex, the ventral temporal cortex, and the hippocampus.

While these studies have highlighted specific facets of human abstract concept generalisation and have employed various modalities to explore it, they have overlooked the temporal dimension which is crucial for certain types of information processing such as those involving auditory objects. Notably, sequences of sounds present such a temporal dimension, where semantic meaning emerges from the temporal concatenation of constituent elements (Bertelson & Tisseyre, 1970; Ladefoged & Broadbent, 1960). Indeed, these sequences can be presented at varying speeds (Höhne & Tangermann, 2012), while the underlying semantic content remains invariant. Previous studies investigating memory for sequences have placed emphasis on how auditory stimulus type and task type modulate recognition and recall.

The literature on auditory sequence perception identifies several key aspects for abstract meaning formation and memorization. First, the length effect: essentially, the more elements a sequence consists of, the more challenging it becomes to remember them to a higher extent (Surprenant et al., 2011; Tehan & Mills, 2007). Schulze & Tillmann (2013) observed a length effect in verbal MW tested by a forward recall task, but not in backward recall. The length effect also modulates performance in recalling pitch sequences of certain characteristics. For example, the pitch proximity effect (higher recall ability for sequences consisting of dissimilar tones than for those made up of similar ones) decreases as the number of items in the sequence increases (Williamson et al., 2010). The tonality effect – tonality enhances recall ability as opposed to atonal sequences – was also found to be countered by increased sequence length, in both forward (Williamson et al., 2010) and backward recall (Croonen, 1994; Schulze et al., 2012). Finally, for retaining sequences of another auditory stimulus type, timbre, reliance on sensory memory was suggested to be the main mechanism (Halpern et al., 2004; Kaernbach, 2004; McKeown et al., 2011).

Although different types of auditory stimuli vary in how challenging it is to remember sequences, they all share the characteristic of having separate components that occur one after the other over time. This temporal aspect of perception aligns well with the principles of Gestalt laws, which offer a framework for understanding how we organise and perceive complex stimuli over time. According to these laws, perception is structured into hierarchical layers, allowing stimuli to be broken down into components without impairing overall recognition (Terhardt, 2016). Thus, even if one characteristic of a stimulus is altered, such as the speed of sound sequences, the holistic perception may remain intact. However, temporal proximity between components can influence this perception, suggesting that the acquired semantic representations might exhibit robustness across different temporal configurations. This robustness could enable the generalisation of learned concepts across varying temporal patterns (Schulze & Tillmann, 2013; Tillmann, 2012).

Despite these studies, the feasibility of such generalisation remains incompletely understood, and the underlying brain mechanisms elusive. To address this question, music emerges as a promising avenue due to its fundamental characteristic of comprising sounds hierarchically organised over time, thereby facilitating the construction of generalisable meaning and, relatedly, a percept of desired musicality in the listener.

For these reasons, particularly when coupled with neuroscientific methods, music has emerged as an excellent medium for investigating auditory perception, memory, information processing, and predictive coding. In our recent studies, we have demonstrated that neural encoding of temporal sound sequences engages a broad network of functionally connected brain areas, particularly in the right hemisphere, including Heschl’s and superior temporal gyri, frontal operculum, cingulate gyrus, insula, basal ganglia, and hippocampus (Bonetti et al., 2021). Similarly, long-term recognition of short musical sequences activates nearly the same brain network as the one engaged during auditory encoding, revealing hierarchical dynamics from lower- to higher-order brain areas during sequence recognition (Bonetti et al., 2023; Fernández-Rubio, Brattico, et al., 2022; Fernández-Rubio, Carlomagno, et al., 2022). Notably, our previous research unveiled faster (150-250 ms from the onset of tones) and negative neural responses to varied musical sequences, contrasting with slower (300-400 ms from the onset of tones) positive activity generated by the brain during the recognition of previously memorised musical sequences (Bonetti et al., 2024).

In the present study, we investigated concept generalisation across different temporal scales using two datasets collected through similar experimental paradigms. Collecting and adding a second set of data, with a different participant pool served for replicating and expanding on the results obtained with the initial dataset. Participants engaged with musical sequences embedded within an old/new recognition paradigm. Specifically, they learned sequences at a defined tempo and were later asked to recognise them and rate their musicality at various tempi, ranging from 150 to 5000 milliseconds of single tone duration, with the original sequence featuring sounds lasting 350 milliseconds. The aim of these experiments was to test four hypotheses.

First, we hypothesised that individuals could generalise abstract concepts across different temporal configurations. Second, even in a temporally generalised context, we expected the brain to distinguish between previously memorised and novel information. Additionally, we anticipated a decline in brain performance with greater distances between learned and generalised items. Third, we predicted that not all brain regions activated during the task would respond differently to various tempi. Some regions would maintain consistent activity across the different sound paces, while others, particularly auditory cortices, might exhibit tempo-dependent responses. Fourth, we expected that lower levels of perceived musicality would be associated with reduced responses in auditory and medial cingulate regions.

## Results

### Experimental design and behavioural results

In Dataset 1, 83 participants completed an old/new auditory recognition task during MEG recording. In this paradigm, first participants were exposed to a short musical piece and asked to memorise it as much as possible (**Figure 1** and **Figure S1**). Second, participants were presented with several five-tone musical sequences and asked to assess whether each sequence belonged to the learned music [‘memorised’ sequence (M), old] or it was a varied sequence [‘novel’ sequence (N), new]. Here, the varied melodies were created by changing all tones of the M sequences from the second one onwards (**Figure S2**). The old/new paradigm was implemented in two subsequent experimental blocks. The first one entailed a learning and a recognition phase, where each tone forming the musical piece and the sequences lasted 350 ms (S1_1_). Here, 27 M and 27 N sequences were presented, in randomised order. Following a 1-minute break in between blocks, the second block consisted only in the recognition phase, since it relied on the same musical piece that was learned in the first block. In this case, to avoid an overly long task, a subset of 21 M and 21 N randomly arranged sequences were presented with two different speeds (S2_1_ = each tone of the sequences lasted 125 ms; S3_1_ = each tone lasted 650 ms, **Figure 1**).

**Figure 1.**
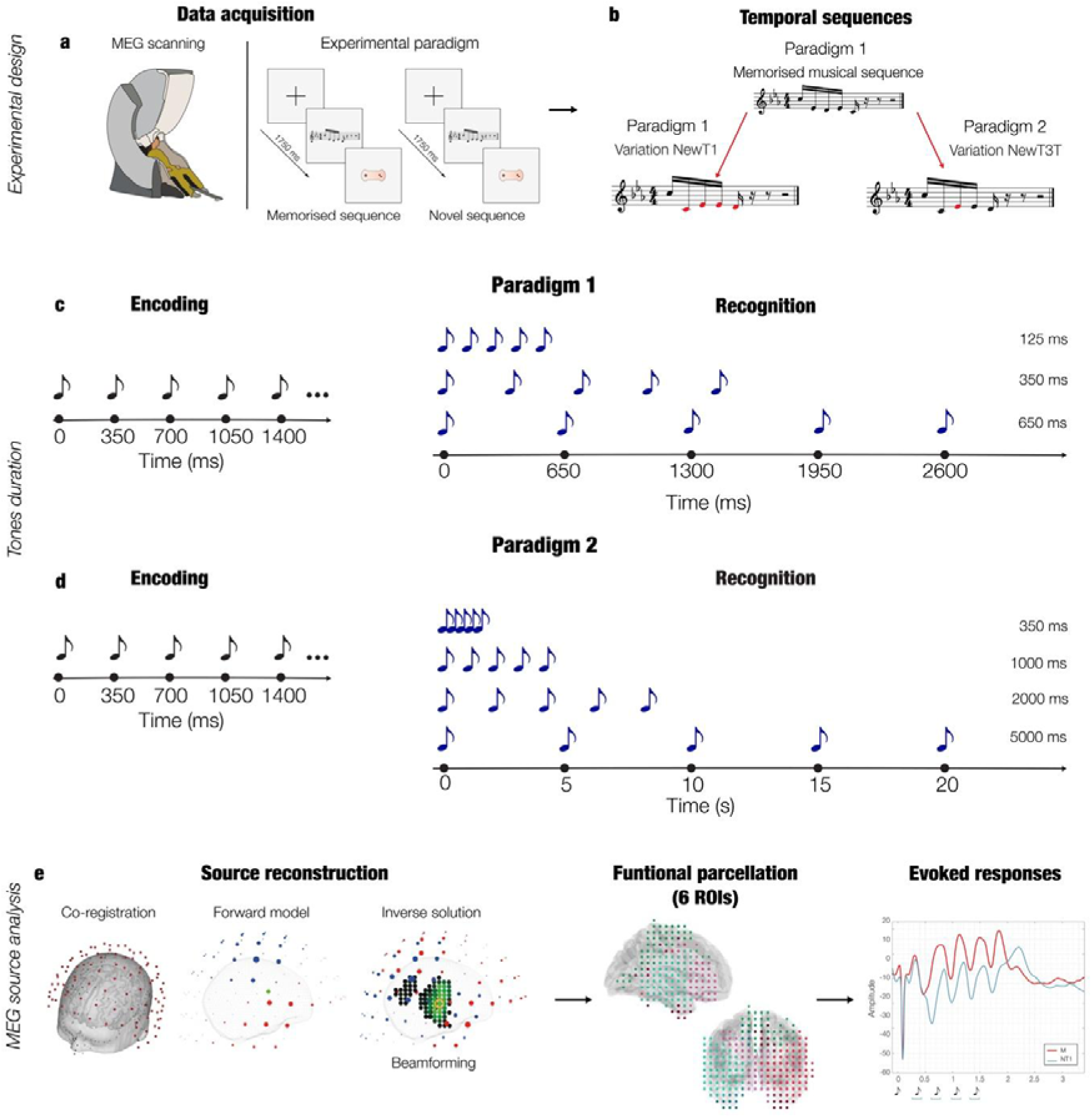
Experimental design, stimuli, and analysis pipeline. **a** – The brain activity of 83 (Dataset 1) and 24 (Dataset 2) participants was collected using magnetoencephalography (MEG) while they performed an auditory old/new recognition task. One at a time, five-tone auditory sequences were presented in randomised order and participants were instructed to respond with button presses whether they were old (memorised musical sequences, M) or new (novel musical sequences, N). **b** – Two types of auditory sequences (M, NT1) were used in the studies (see **Figure S1** for the prelude and **Figure S2** for the full set of sequences in musical notation). The N sequences of paradigm 1 were created through systematic variations (V.) of the M sequences. This procedure consisted of changing every musical tone of the sequence after the first (NT1) tone, as illustrated by the red musical tones. The N sequences of paradigm 2 were created by systematically changing only the third tone **c** – The experimental procedure of Dataset 1 consisted first of an encoding phase, where the length of notes forming the musical piece to be learned lasted 350 ms. Second, the M and NT1 sequences described in **b** were presented at three different speeds: each tone lasting either 350 ms, 125 ms or 650 ms. **d –** The experimental paradigm of Dataset 2 also consisted of an encoding and a recognition phase. The former was the same as for the paradigm described in **c**. The latter consisted of presenting M and NT3T sequences at four different speeds: each tone lasting either 350 ms, 125 ms or 650 ms. **e –** The MEG data was co-registered with the individual anatomical magnetic resonance imaging (MRI) data, and source reconstructed using a beamforming algorithm. This resulted in one time series for the 3559 reconstructed brain sources (8-mm parcellation) which were then conveyed into six functional regions of interest (ROIs), as described in Bonetti et al. (2024). Finally, the reconstructed brain activity was contrasted across experimental conditions (M versus N), independently for each time-point, ROI and speed and corrected for multiple comparisons using one-dimensional (1D) cluster-based Monte-Carlo simulations (MCS).

In Dataset 2, 24 participants completed the same old/new paradigm described for Dataset 1. Here, the learning occurred using the same musical prelude described for Dataset 1, where each tone lasted 350 ms. Conversely, for the recognition phase, the varied sequences were constructed by changing only the 3^rd^ tone of the original M sequences (and not all the sounds after the 1^st^ one). The duration of each tone was also different, comprising the following speeds: S1_2=_ each tone of the sequence lasted 350 ms; S2_2_ = 1000 ms; S3_2_ = 2000 ms; S4_2_ = 5000 ms (**Figure 1**). Moreover, after performing the memory recognition task, participants rated the perceived musicality of the M sequences over the different speeds.

All MEG task behavioural data was analysed computing independent Kruskal-Wallis H tests, in order to reveal potential differences in response accuracy between experimental conditions, independently for Dataset 1 and Dataset 2. Additionally, perceived musicality ratings from Dataset 2 were also analysed using the same statistical methods (**Figure 2**).

**Figure 2.**
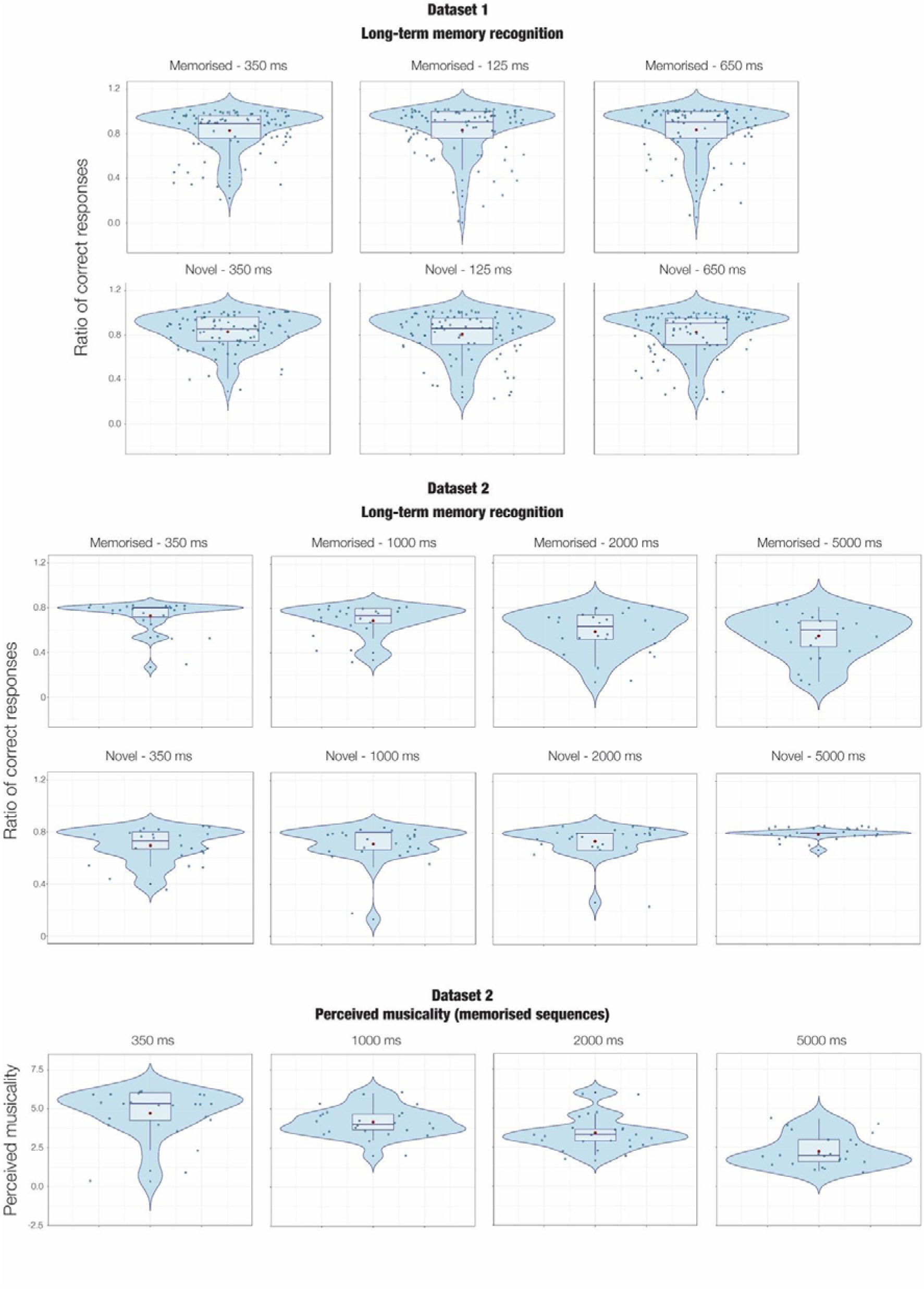
Behavioural performance in the MEG auditory recognition task. Scatter, violin and box plots overlaid to illustrate the number of correct responses for each experimental task of Dataset 1 and Dataset 2. The plots depict each experimental condition separately. Each dot represents a participant (Dataset 1 - n = 83; Dataset 2 - n = 24). The box plots depict the median, the interquartile range, while the lines extending from the box show the range of non-outlier data points. The violin plots visualise the distribution of the data along the y-axis. Here, the red dot illustrates either the mean response accuracy for the old/new recognition paradigms, or the mean ratings for perceived musicality. The Tukey-Kramer correction following the Kruskal-Wallis H test has displayed no significant behavioural level result for Dataset 1, while for Dataset 2 the following pairs differ significantly from each other in terms of response accuracy: M 350 vs M 2000, p = .003; M 350 vs M 5000, p = .001; N 350 vs. M 2000, p = .002; N 350 vs. M 5000, p < .001; M 1000 vs. N 5000, p = .007; N 1000 vs. M 2000, p < .001; N 1000 vs. M 5000, p < .001; M 2000 vs. N 2000, p < .001; M 2000 vs. 5000 N, p < .001; N 2000 vs. M 5000, p < .001; M 5000 vs. N 5000, p < .001, and the following in terms of perceived musicality: M 350 vs. M 2000, p = .005; M 350 vs. M 5000, p < .001; M 1000 vs. 5000, p < .001.

The Kruskal-Wallis H test contrasting response accuracies of M and N conditions, across all speeds of Dataset 1 revealed no significant results (*p* = .32).

For the recognition task of Dataset 2, the Kruskal-Wallis H test contrasting response accuracy across all conditions and speeds was significant (H(7) = 73.49, *p* = 2.90e-13), pointing to divergence between categories in number of accurate responses. The Tukey-Kramer correction for multiple comparisons showed the following significant comparisons: M 350 vs M 2000, *p =* .003; M 350 vs M 5000, *p =* .001; N 350 vs. M 2000, *p* = .002; N 350 vs. M 5000, *p* < .001; M 1000 vs. N 5000, *p* = .007; N 1000 vs. M 2000, *p* < .001; N 1000 vs. M 5000, *p* < .001; M 2000 vs. N 2000, *p* < .001; M 2000 vs. 5000 N, *p* < .001; N 2000 vs. M 5000, *p* < .001; M 5000 vs. N 5000, *p* < .001.

Third, perceived musicality ratings were also examined with a Kruskal-Wallis H test, that was significant (H(7) = 39.22, *p* = 1.56e-08). The Tukey-Kramer correction for multiple comparisons highlighted a significant difference between musicality ratings for the following pairs: M 350 vs. M 2000, *p* = .005; M 350 vs. M 5000, *p* < .001; M 1000 vs. 5000, *p* < .001. This indicates that lower perceived musicality was associated with the slower speeds.

**Figure 2** illustrates the behavioural analyses, revealing a decline in performance in recognising novel sequences as the duration of the tones increased. Similarly, perceived musicality deteriorated with longer durations of the musical tones.

### Neural sources

The MEG sensor data was pre-processed using standard procedures and its neural sources reconstructed using a beamforming approach (**Figure 1**). This procedure sequentially applies a different set of weights to the source locations, in order to identify the contribution of each source to the activity recorded by the MEG sensors. This is a widely adopted procedure which is described in detail in the Methods section. The beamforming algorithm was employed in conjunction with an 8-mm parcellation of the brain, resulting in 3559 distinct brain sources, each characterised by its own time series. In this study, we were specifically interested in the brain regions of interest (ROIs) primarily recruited during musical recognition memory processes. Thus, we selected the six large ROIs described by Bonetti et al. (2024) who used the same paradigm but in a different context (shown in **Figure S3**). Here, the time series of the brain voxels within each of the six ROIs were averaged, obtaining only one time series for each ROI. Then, we computed one two-sided t-test contrasting M versus N conditions, independently for each ROI, each time-point and speed. To correct for multiple comparisons, we used one-dimensional (1D) cluster-based Monte-Carlo simulations (MCS, α = .05, MCS *p-value* = .001). The same procedure was computed independently for the two Datasets.

In essence, results (**Figures 3, 4** and **5**) revealed several clusters of significant differential activity between M and N across the selected ROIs and the speeds included in the study. More specifically, the contrast M versus N elicited stronger activity in RHG, LHP, RHP and ACC at 350 – 450 ms after the onset of each tone. Additionally, M was characterised by a stronger negativity than N in the MC at 400-500 ms post-onset after each tone (**Figure 5**). Conversely, LHP, RHP and VMPFC demonstrated a stronger response for N versus M observable at 250 – 300 ms after the tones which modified the original sequences (**Figures 4** and **5**). The magnitude of the differential activity between M and N was reduced when the musical sequences were presented at a slower speed. Detailed information on the largest significant clusters is reported in **Table 1**, while complete statistical results are extensively reported in **Table S1.**

**Figure 3.**
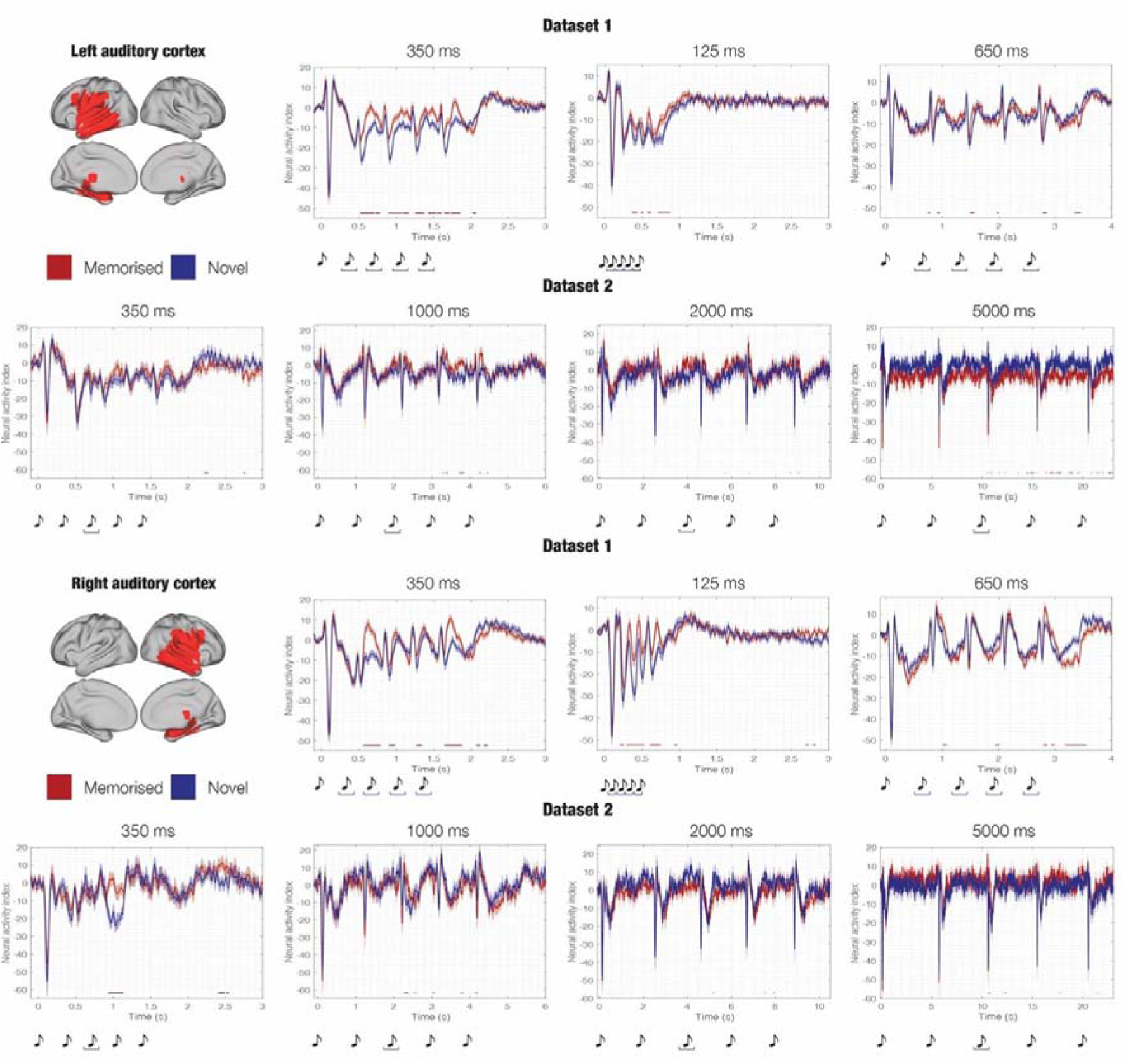
Source-localised differences in evoked responses across experimental conditions. This figure illustrates the source-localised brain activity averaged over participants (Dataset 1 - n = 83; Dataset 2 – n = 24) for each experimental condition of Dataset 1 and Dataset 2 (memorised [M], novel T1 [NT1] and memorised [M], novel T3T [NT3T]) within two selected automated anatomical labelling (AAL) regions of interest (ROIs): left auditory cortex (ACL) and right auditory cortex (ACR). Brain templates depict the spatial extent of the selected ROIs. Here, the contrasts were computed using two-sided t-tests, independently for each time-point, and correcting for multiple comparisons using one-dimensional Monte-Carlo simulations (MCS; MCS, α = .05, MCS p-value = .001). Red and blue correspond to specific M versus N condition comparisons, while the musical tones depict the onset of each sound forming the sequences. For a comprehensive depiction of all AAL ROIs, please refer to **Figure S3**. For a detailed statistical report on significant differences between experimental conditions in all ROIs, consult **Table S1**.

**Figure 4.**
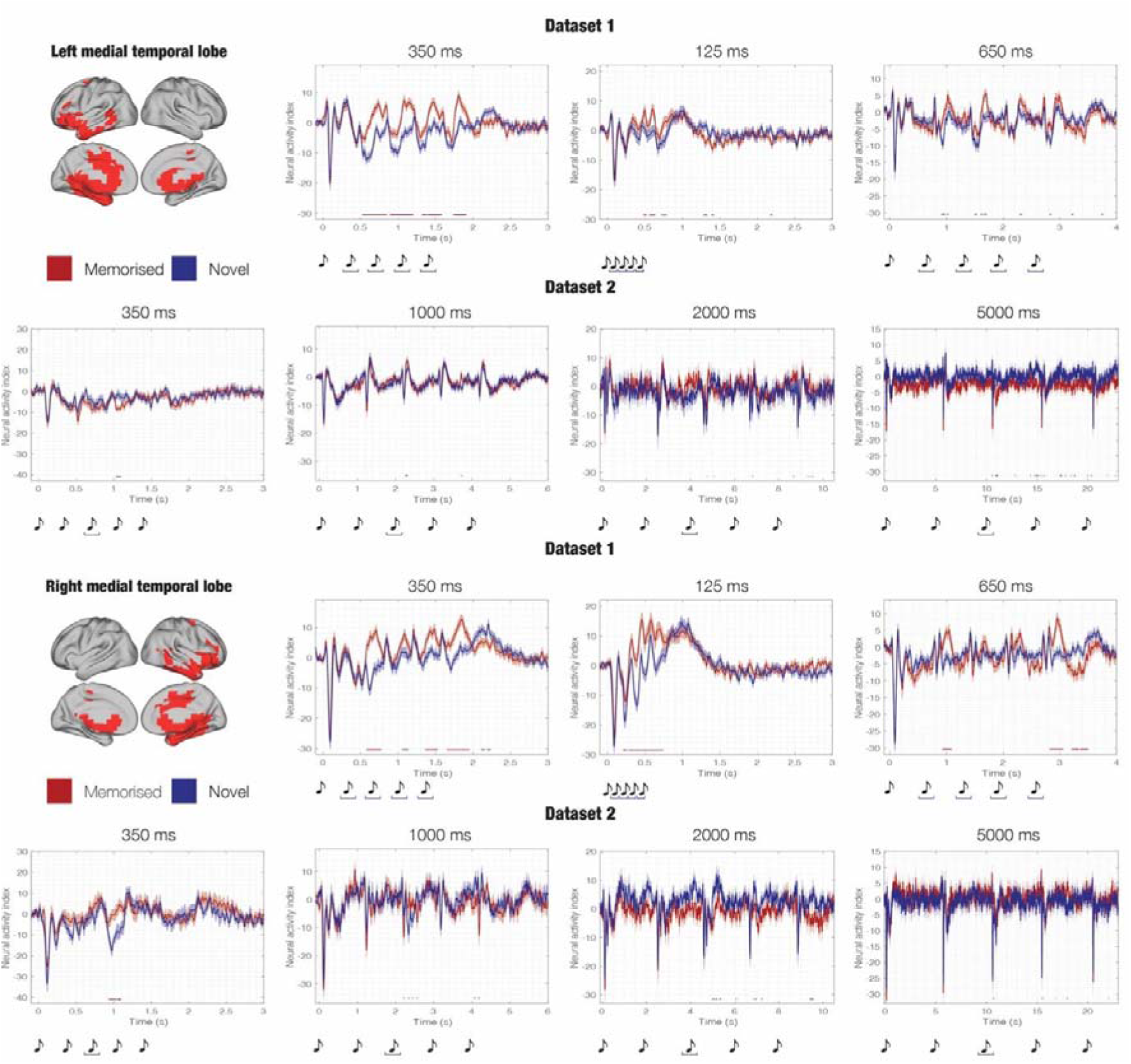
Source-localised differences in evoked responses across experimental conditions. This figure illustrates the source-localised brain activity averaged over participants (Dataset 1 - n = 83; Dataset 2 – n = 24) for each experimental condition of Dataset 1 and Dataset 2 (memorised [M], novel T1 [NT1] and memorised [M], novel T3T [NT3T]) within two selected automated anatomical labelling (AAL) regions of interest (ROIs): left medial temporal lobe (MTLL) and right medial temporal lobe (MTLR). Brain templates depict the spatial extent of the selected ROIs. Here, the contrasts were computed using two-sided t-tests, independently for each time-point, and correcting for multiple comparisons using one-dimensional Monte-Carlo simulations (MCS; MCS, α = .05, MCS p-value = .001). Red and blue correspond to specific M versus N condition comparisons, while the musical tones depict the onset of each sound forming the sequences. For a comprehensive depiction of all AAL ROIs, please refer to **Figure S3**. For a detailed statistical report on significant differences between experimental conditions in all ROIs, consult **Table S1**.

**Figure 5.**
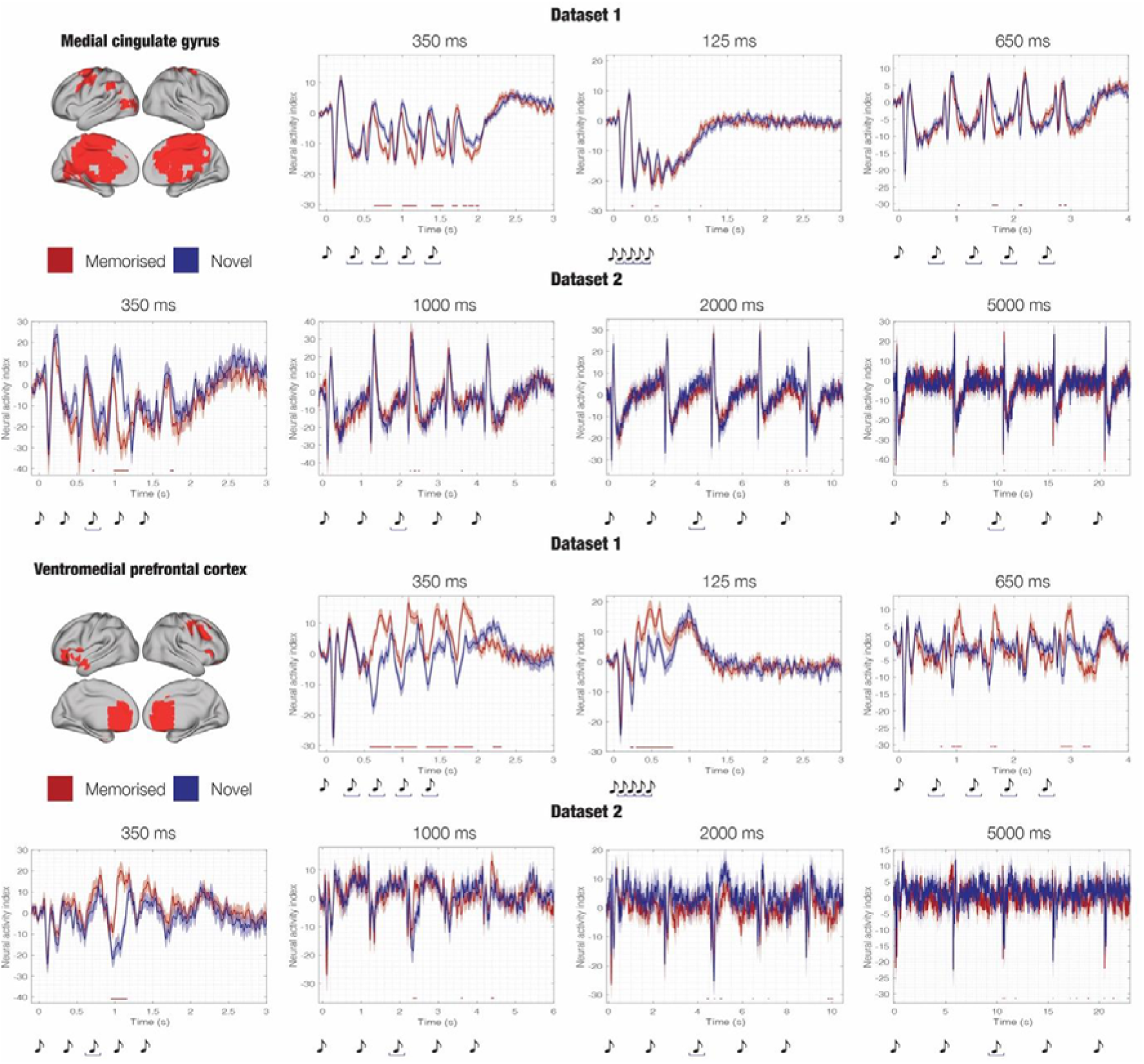
Source-localised differences in evoked responses across experimental conditions. This figure illustrates the source-localised brain activity averaged over participants (Dataset 1 - n = 83; Dataset 2 – n = 24) for each experimental condition of Dataset 1 and Dataset 2 (memorised [M], novel T1 [NT1] and memorised [M], novel [NT3T]) within two selected automated anatomical labelling (AAL) regions of interest (ROIs): medial cingulate gyrus (MCG) and ventromedial prefrontal cortex (VMPFC). Brain templates depict the spatial extent of the selected ROIs. Here, the contrasts were computed using two-sided t-tests, independently for each time-point, and correcting for multiple comparisons using one-dimensional Monte-Carlo simulations (MCS; MCS, α = .05, MCS p-value = .001). Red and blue correspond to specific M versus N condition comparisons, while the musical tones depict the onset of each sound forming the sequences. For a comprehensive depiction of all AAL ROIs, please refer to **Figure S3**. For a detailed statistical report on significant differences between experimental conditions in all ROIs, consult **Table S1**.

**Table 1.** Largest clusters of stronger activity of memorised (M) versus novel sequences (N). Largest clusters of significantly stronger activity of M versus N computed for the six regions of interest (ROIs): left auditory cortex (ACL), right auditory cortex (ACR), left medial temporal lobe and inferior temporal cortex (LMTL), right medial temporal lobe and inferior temporal cortex (RMTL), medial cingulate gyrus (MC), and ventromedial prefrontal cortex (VMPFC). The table depicts the contrast (two-sided t-tests), the temporal extent of the largest significant cluster (ms), the associated ROI, the peak t-value of the cluster and the corresponding Monte-Carlo simulations (MCS) p-value. This information is reported independently for the two Datasets of the study. Extensive statistical details are available in **Table S1**.

| *Stimulus speed* | ROI | Temporal extent of the largest clusters from the 1<sup>st</sup> tone of the sequence (ms) | Peak t-value | P-value |
| --- | --- | --- | --- | --- |
| *Dataset 1* |  |  |  |  |
| 125 ms | ACL | 708 – 872 | 4.613 | < .001 |
|  | ACR | 296 – 532 | 7.511 | < .001 |
|  | LMTL | 716 – 792 | 3.457 | < .001 |
|  | RMTL | 280 – 744 | 7.846 | < .001 |
|  | MCG | 540 – 592 | -3.092 | < .001 |
|  | VMPFC | 292 – 780 | 5.918 | < .001 |
| 350 ms | ACL | 892 – 1084 | 5.872 | < .001 |
|  | ACR | 556 – 792 | 7.798 | < .001 |
|  | LMTL | 524 – 852 | 6.148 | < .001 |
|  | RMTL | 1648 – 1952 | 5.454 | < .001 |
|  | MCG | 628 – 860 | -5.240 | < .001 |
|  | VMPFC | 896 – 1192 | 5.498 | < .001 |
| 650 ms | ACL | 3340 – 3452 | -4.403 | < .001 |
|  | ACR | 3168 – 3556 | -5.271 | < .001 |
|  | LMTL | 1648 – 1716 | 3.507 | < .001 |
|  | RMTL | 2816 – 3056 | 4.974 | < .001 |
|  | MCG | 1620 – 1724 | -4.210 | < .001 |
|  | VMPFC | 2816 – 3024 | 4.021 | < .001 |
| *Dataset 2* |  |  |  |  |
| 350 ms | ACL | 2220 – 2272 | -2.798 | < .001 |
|  | ACR | 936 – 1136 | 5.121 | < .001 |
|  | LMTL | 1032 – 1100 | -3.063 | < .001 |
|  | RMTL | 936 – 1040 | 4.477 | < .001 |
|  | MCG | 984 – 1180 | -6.355 | < .001 |
|  | VMPFC | 948 – 1164 | 6.890 | < .001 |
| 1000 ms | ACL | 4252 – 4296 | 3.521 | < .001 |
|  | ACR | 2288 – 2368 | 3.123 | < .001 |
|  | LMTL | 3712 – 3740 | 2.894 | < .001 |
|  | RMTL | 2320 – 2364 | 3.243 | < .001 |
|  | MCG | 2360 – 2424 | -2.666 | < .001 |
|  | VMPFC | 2336 – 2440 | 4.749 | < .001 |
| 2000 ms | ACL | 6784 – 6812 | 3.186 | < .001 |
|  | ACR | 5152 – 5184 | -2.981 | < .001 |
|  | LMTL | 4740 – 4772 | 3.667 | < .001 |
|  | RMTL | 4984 – 5084 | -3.822 | < .001 |
|  | MCG | 8208 – 8252 | -3.564 | < .001 |
|  | VMPFC | 5036 – 5080 | -2.537 | < .001 |
| 5000 ms | ACL | 14 984 – 15 152 | -4.043 | < .001 |
|  | ACR | 10 624 – 10 764 | 4.110 | < .001 |
|  | LMTL | 15 016 – 15 168 | -4.550 | < .001 |
|  | RMTL | 10 716 – 10 764 | 3.274 | < .001 |
|  | MCG | 20 560 – 20 604 | 3.638 | < .001 |
|  | VMPFC | 18 956 – 18 988 | -3.515 | < .001 |

## Discussion

This study investigates the generalisation of the recognition of abstract concepts, specifically those with a temporal component. To enhance our understanding, we compared two datasets (see Methods and Results for detailed descriptions) using an old/new auditory recognition paradigm. In this paradigm, while their neural activity was recorded using MEG, participants were presented with musical sequences at varying speeds and were asked to determine whether each sequence was one they had learnt before the task or was a variation. Building on our previous research on long-term memory for musical sequences (Bonetti, Brattico, et al., 2023; Bonetti et al., 2020; Bonetti, Fernández-Rubio, et al., 2023; Fernández-Rubio, Brattico, et al., 2022; Fernández-Rubio, Carlomagno, et al., 2022; Fernández-Rubio et al., 2024) this study incorporates a temporal dimension into the material learnt in the old/new recognition paradigm. While continuing to examine auditory sequence recognition, this study also investigates in-depth the ability and neural mechanisms of abstract concept generalisation and meaning formation, through the alteration of stimulus speed in both memorised and novel sequences in association with musicality ratings. The results from the two datasets presented in this study provide insights into the encoding of the overarching meaning of a musical sequence and how such meaning can be generalised to variations in their temporal configuration. The findings demonstrate the ability to generalise such concepts, both on the behavioural and neural levels.

In line with our previous studies, both datasets showed an average response accuracy of at least 70%, indicating a highly refined ability to distinguish between old and new auditory sequences. Interestingly, while Dataset 1 presented only relatively small differences in the speeds of the sequences, resulting in no changes in the participants’ performance, Dataset 2 involved comparatively larger-scale differences in stimulus duration. As expected, the decline in performance observed for the N sequences was proportional to the increase in their duration. This trend is partially in accordance with the results of Dowling et al., (2008), who have discovered a detrimental effect on the recognition of familiar melodies of both speeding up and slowing down the tempo.

Our findings are partially corroborated by the work of Schulze & Tillmann (2013), who explored recognition memory for pitch sequences. They identified a "length effect," where WM performance declined as the number of items in the sequence increased. Unlike timbre sequences, pitch sequences can be rehearsed, which generally enhances recognition. This comparison highlights that, although both fall under the same sensory modality, different stimulus components uniquely influence encoding. These insights provide a clearer understanding of our own results. In our study, we focused on pitch sequences. However, we did not modulate the length of the sequences but the speed of their presentation. Similarly to their study, we found that the recognition ability decreased as a function of the length of the presented sequences, possibly due to a limited capacity of the human brain to retain the representation of the sounds. Interestingly, such relation only involves the N sequences and does not relate to the M melodies. A possible explanation for that is that, as the speed increases, the differences between the two conditions become more pronounced, with the proportion of hits consistently being lower for N sequences at faster speeds. In other words, comparing the recognition of M and N sequences under optimal conditions likely shows no significant difference. However, when experimental conditions are modified (e.g., decreased speed), differences in recognition performance between the M and N conditions become more pronounced. These findings align with previous research by Fernández-Rubio et al. (2024), which compared recognition memory between older and younger adults. In that study, older adults performed worse than younger adults when recognising novel melodies, but there was no significant difference in detecting previously memorised musical sequences. Thus, cognitive decline associated with aging impairs the recognition of N melodies but not previously memorised ones (M), suggesting that recognition of N melodies is more complicated than M, coherently with the results presented in the current study.

In line with our previous findings (Bonetti et al., 2024), our study reinforces the applicability of predictive coding theory (PCT) in understanding auditory processing mechanisms (Millidge et al., 2022). While our results also align with other theories such as Bayesian Brain Theory and Hierarchical Temporal Memory, PCT offers a more comprehensive explanation for the observed phenomena. Specifically, the prevalent prediction error observed across regions such as the medial temporal lobe and ventromedial prefrontal cortex can be attributed to PCT, as it accounts for the disruption of top-down predictions by incoming sensory stimuli. In our study, this prediction error is manifested as a negative component peaking around 250 ms post stimulus-onset, reflecting the discrepancy between the expected and heard tones. Additionally, we observed strong right lateralisation in response strength, indicative of robust neural processing. Notably, the consistency of prediction error across varying stimulus speeds, particularly for sounds shorter than 1000 ms, underscores the concept of generalisation in hierarchical processing within the brain. This suggests that the mechanisms underlying auditory prediction may extend across different temporal scales, contributing to a deeper understanding of cognitive processing.

In our study, we also asked participants to rate the musicality of the memorized sequences of Dataset 2, as a proxy for meaning formation of a generalized concept, namely a melody (being it played at a slower or a faster pace). The results unanimously indicate that stimuli presented at slower speeds were perceived as less musical. Given that recognition performance for N melodies declined at slower speeds, one possible explanation is that the large time intervals between stimuli made chunking no longer possible. A previous behavioral study (Warren et al. 1991) showed similar effects but with highly familiary melodies and durations ranging between 40 and 5000 milliseconds and concluded that stimuli presented "too fast" negatively impacted both recognition and perceived musicality. These findings provide valuable insight into the process of making sense of musical sounds, by highlighting how the efficient temporal integration of items into episodic memory is functional not only to allow a successful recognition of musical motifs but also to produce a positive aesthetic experience in the listener. Sound events that occur too fast might impair the decision-making process of attribution of a subjective, abstract meaning to melodies, which is well captured by holistic ratings of musicality. Such interpretation resonates with the predictive processing account of aesthetic reactions to music, according to which aesthetic pleasure originates when prediction of incoming sounds can be efficiently implemented. When predictive models are too precise or too obscure, the listener does not achieve a positive aesthetic experience whereas when the incoming sound does not match the model and this model deviation can be smoothly resolved by model updating and active sensorimotor regulation then a positive aesthetic response is issued (Brattico & Delussi, 2024; Vuust et al., 2022), and this process seems to occur not only in music but also in other forms of art, due to the evolutionarily positively valued experience of resolving the uncertainties in the environment (Van de Cruys & Wagemans, 2011).

Overall, the neural results showed several significant clusters of differences between the experimental conditions, with effect sizes varying as a function of the speed of the melody presentation. First, the neural response patterns of the six ROIs at the baseline speed of 350 ms further reinforce the results reported in Bonetti et al. (2024). Second, the neural results exhibited a coherent trend compared to the behavioural results. Specifically, the strength of the difference between M and N conditions decreased as the duration of the stimuli increased (i.e., as speed decreased). One possible explanation for this pattern is reminiscent of the Gestalt principle of continuity, as stimulus continuity - especially over durations longer than 1000 ms - is disrupted, making perception more difficult (Sutojo et al., 2020). In contrast, higher stimulus speeds can be perceived as holistic patterns, thereby facilitating recognition.

The aspect of a holistic pattern may also be relevant when examining the activity of the medial cingulate gyrus (MC). This region exhibits activity for about 1000 ms following each tone before returning to baseline. This pattern is clearly visible in all speeds, and especially in the cases of the 2000 ms and 5000 ms duration of each sound. The systematic return to baseline could suggest a role in connecting the items of the sequences, which are, in this case, the individual tones. When this connectivity is disrupted, recognition worsens, and perceived musicality decreases, pointing towards a possible role of MC in providing the holistic perception which humans experience when listening to music, as opposed to single sounds. This claim is supported by our previous research linking the cingulate gyrus to musical perception and musical expertise (Criscuolo et al., 2022; Pando-Naude et al., 2021) as well as by the work of Koelsch (2018, 2020).

Examining the signals elicited by the N sequences, it is noticeable that the prediction error to the varied sounds occurs across all stimulus durations, showing the typical negative peak around 250-300 ms after the onset of the varied sounds. However, the strength of this response declines as stimulus duration increases, with a noticeable cutoff at 1000 ms between-tone intervals. At this point, the negative response appears to be replaced by a very late positive component peaking approximately 1000 ms after the varied sounds. Moreover, the neural responses to the stimuli of Dataset 2 reveal an interesting tendency, highlighting the value of the paradigm differences between the two datasets. As expected, prediction error occurs, elicited by the third, altered tone (at all speeds), and only here is the difference between conditions significant. Unlike in Dataset 1, this is the only tone altered in the sequence. However, the response strength following the fourth and fifth tones (those after the altered tone) matches that given to the same tones in the memorised sequence. This implies that, in addition to the sequence being encoded and remembered as a chunk conveying an overarching meaning, the individual tones are also encoded and monitored by the brain.

Previous neuroimaging studies investigating concept generalisation have identified key brain regions that partially overlap with the findings presented in our current study. For instance, Bowman & Zeithamova, (2018) focused on abstract visual concepts and identified the ventromedial prefrontal cortex (VMPFC) and the hippocampus as crucial for generalisation. While these regions align with our own findings, discrepancies in lower-level structures may arise due to the different sensory modalities employed. Sheikh et al. (2021) explored concept generalisation across languages, identifying several regions including the VMPFC, temporal lobe structures, the posterior cingulate gyrus, and parts of the parietal lobe. While our study complements these findings, further research is warranted to fully elucidate the neural mechanisms underlying concept generalisation in the auditory domain, considering the diverse sensory and cognitive processes involved.

The recorded responses in the brain exhibit distinct patterns depending on the speed of the stimuli. Specifically, the auditory cortices and the medial cingulate gyrus demonstrate response patterns that appear to be more sluggish when stimuli are presented at slower speeds, particularly until the duration reaches no longer than 1000 ms for each sound. In contrast, the medial temporal lobes and the ventromedial prefrontal cortex maintain relatively consistent activity levels even when the duration of stimuli is manipulated. This duality in observed responses suggests that regions exhibiting consistent activity independent of speed may play a crucial role in the generalisation ability demonstrated in the study. By maintaining stable responses across varying speeds, these regions may contribute to the brain’s capacity to generalise auditory information and adapt to dynamic environmental stimuli.

Lerner et al. (2014) investigated whether neural response rates are adjusted to the speed of presented stimuli in the structures building up the network involved in speech processing. When analysing data of speech comprehension rates over different speeds, a positive effect of speeding up the stimuli was described, both in terms of individual word recognition and sentence recognition, whereas stimuli over prolonged durations were harder to extract meaning from. The fMRI data collected under the scope of the same study has revealed that temporal scaling of responses depending on the speed the stimulus is presented with was prevalent in case of the sensory areas, in case the auditory cortices, whereas this tendency diminished in case of higher-order brain regions.

While the present study yields strong, consistent, and clear results, several limitations warrant consideration. Firstly, Dataset 2 suffers from a relatively small sample size and limited trial numbers, which may impact the generalisability of findings. Additionally, the slight divergence in paradigms between the two datasets - specifically, the variation in the timing of stimulus alterations - hinders direct comparability. Moreover, the restriction of interval ranges to above 150 ms poses a limitation on the study’s ability to capture responses to shorter temporal intervals due to overlapping responses and participant discernibility issues, as noted by Warren et al. (1991). Despite these constraints, the robustness of our results remains evident.

Moving forward, future research endeavours could explore the generalisation of non-musical temporal stimuli, such as sequences of numbers, to broaden our understanding of temporal concept generalisation. Furthermore, investigating sequences of visual elements could elucidate whether the brain network identified in the present study is universally involved in different types of temporal concept generalisation or selectively music related. By addressing these avenues, we can further refine our understanding of the neural mechanisms underlying concept generalisation across diverse domains.

## Materials and methods

### Participants – Dataset 1

The data was collected from a sample of 83 participants [50 females and 33 males (sex, biological attribute, self-reported)] aged 19 to 63 years old (mean age: 28.76 ± 8.06 years). All participants came from Western countries, and the data acquisition took place in Aarhus, Denmark. All participants in the sample reported normal hearing and were drawn from a generally homogeneous and highly educated population, with most participants with at least one university degree. The Institutional Review Board (IRB) of Aarhus University has approved the project (case number: DNC-IRB-2020-006). All experimental procedures were complying with the Declaration of Helsinki – Ethical Principles for Medical Research. All participants have given their informed consent prior to partaking in the experimental procedure.

### Experimental stimuli and design – Dataset 1

The experimental procedure consisted of magnetoencephalography (MEG) recordings of participants’ brain activity while they completed an old/new paradigm, developed in the context of auditory long-term recognition. The procedure was divided into two blocks, carried out during the same recording session. The first block entailed a learning phase, where participants were asked to memorise a short musical piece, which was played for them twice, and a recognition phase, in which participants were presented with several melodies and required to distinguish between previously memorised (excerpts from the short musical piece) and novel sequences. The musical piece to be memorised in the learning phase was taken from Johann Sebastian Bach’s Prelude No. 2 in C Minor, BWV 847. More precisely, as shown in **Figure S1**, it was the first four bars of the right-hand part, where each bar included 16 tones, meaning that the total number of tones added up to 64 (16*4). The duration of each tone was approximately 350 ms, for 22400 ms in total. Furthermore, a final tone - lasting for 1000 ms - was included following the four bars, in order to provide musical closure. Thus, the whole duration was 23400 ms (23.4 seconds). Finale (MakeMusic, Boulder, CO) was used to create a MIDI version of the piece. In the recognition phase, 135 five-tone musical excerpts were played for the participants, each lasting 1750 ms. The participants’ task was to categorise each excerpt, based on whether it belonged to the original music (‘memorised’ sequence [M], old), or was a modified musical sequence (‘novel’ sequence [N], new) (**Figure 1**). The first block used the same stimuli reported in Bonetti et al., (2024) where M and N were constructed as follows: the M sequences consisted of the first five tones of the first three measures of the musical piece, resulting in three different memorised sequences, each repeated nine times, for a total of 27 M sequences. The N sequences were the result of changing every musical tone of the melody after the first (NT1), second (NT2), third (NT3) or fourth (NT4) tone. Each category of N sequences was composed of 27 sequences. While Bonetti et al., (2024) reported the results for all stimuli, NT2, NT3, NT4 did not serve the scope of the current study and thus were not included in the analyses described below. Additional details on the creation of the sequences can be found in Bonetti et al., (2024).

In the second block, only the recognition phase was implemented. Here, the speed of the M and NT1 sequences was varied. While the duration of each tone of the sequences in block 1 lasted 350 ms, here two new categories of M and NT1 were created, where each sound lasted either 125 ms or 650 ms. To be noted, the M and NT1 sequences consisted of the exact same tones as the ones memorised during the encoding phase of block 1, only over different speeds. Here, participants performed the same old/new recognition task described for block 1, relying on the musical information memorised in the first learning phase of block 1. In this recognition task, a total of 84 trials were presented to the participants, in a randomised order, with 21 sequences from each category (M – 125 ms, N – 125 ms, M – 650 ms, N – 650 ms). The 21 M and 21 NT1 stimuli constituted subsets of the 27 M and NT1 stimuli described for the first experimental block, in order to keep the experiment short. All musical sequences are available in **Figure S2**.

### Participants – Dataset 2

The participant pool consisted of 24 people (8 males and 16 females) voluntarily partaking in the experimental procedure. They had all given their informed consent before following through with the experiment. They were between the ages of 19 and 51 years old (mean age: 28.17 ± 8.58 years) and have reported normal hearing. The participants shared homogeneous and highly educated backgrounds, with the majority holding at least one university degree. The collection of the data took place in the Aarhus University Hospital (AUH), Aarhus, Denmark. The Institutional Review Board (IRB) of Aarhus University has approved the project (case number: DNC-IRB-2020-006). All experimental procedures were complying with the Declaration of Helsinki – Ethical Principles for Medical Research.

### Experimental stimuli and design – Dataset 2

Dataset 2 consisted of exposing the participants to the same auditory old/new paradigm presented for Dataset 1, which entailed a learning and a recognition phase. The Bach’s musical piece played during the learning phase was identical to the one used in Dataset 1, as described above. In the recognition phase, 108 five-tone musical sequences were played for the participants. Exactly as for dataset 1, the M sequences consisted of the first five tones of the first three measures of the musical piece presented in the learning phase, resulting in three different memorised sequences. Here, each of them was played at four different speeds (S1 = each tone lasted 350 ms, S2 = 1000 ms, S3 = 2000 ms, S4 = 5000 ms) and repeated four times, for a total of 12 M sequences for each speed (48 in total). The N sequences were instead constructed in a slightly different way compared to Dataset 1. Here, only the tone number three of the sequences was changed and thus the N sequences were named NT3T. Five N sequences were created for each speed and each of the three M sequences, for a total of 60 N sequences. As for Dataset 1, participants were instructed to categorise each of the sequences as M or N. **Figure S2** depicts all the sequences used for Dataset 2.

Finally, after finishing the main memory task, participants were asked to rate the M sequences played in the different speeds in terms of perceived musicality, using a scale ranging between 1 (minimal musicality) and 7 (extreme musicality).

### Data acquisition

The following paragraphs concerning data acquisition apply to both datasets.

The MEG data was acquired using an Elekta Neuromag TRIUX MEG scanner with 306 channels (Elekta Neuromag, Helsinki, Finland), in a magnetically shielded room at Aarhus University Hospital (AUH), Denmark. The data was recorded at a sampling rate of 1000 Hz with an analogue filtering of 0.1 – 330 Hz. The head shape of the participants and the position of four Head Position Indicator (HPI) coils were registered according to three anatomical landmarks using a 3D digitizer (Polhemus Fastrak, Colchester, VT, USA) prior to the MEG recordings. This data was utilised to co-register the MEG-recordings with the anatomical MRI scans. Throughout the MEG recordings, the continuous localisation of the head was registered by the HPI coils. This was employed for movement correction afterwards. Additionally, cardiac rhythm and eye movements were recorded using two sets of bipolar electrodes, allowing for the removal of electrocardiography (ECG) and electrooculography (EOG) artifacts during later stages of the analysis workflow.

The MRI scans were recorded at AUH, using a CT-approved 3T Siemens MRI-Scanner. The acquired data entailed structural T1 (mprage with fat saturation) with a spatial resolution of 1.0 x 1.0 x 1.0 mm and the following sequence parameters: echo time (TE) = 2.61 ms, repetition time (TR) = 2300 ms, reconstructed matrix size = 256 x 256, echo spacing = 7.6 ms, bandwidth = 290 Hz/Px.

The MEG and MRI data were acquired two separate days.

### Behavioural data

Behavioural data was obtained from the experimental task participants completed during the MEG recording for the two Datasets. This data entails the number of correctly recognised musical sequences. Since the data was not normally distributed, one independent Kruskal-Wallis H test (non-parametric one-way analysis of variance) was computed for each dataset to determine whether the correct responses in the recognition task were different between M and N and over the various speeds. Tukey-Kramer correction was used to correct for multiple comparisons. The perceived musicality ratings collected in Dataset 2 were also evaluated using a Kruskal-Wallis H test, followed by Tukey-Kramer correction.

### MEG data pre-processing

The two datasets were pre-processed in the exact same fashion. The first pre-processing of the raw MEG sensor data (204 planar gradiometers and 102 magnetometers) was done by MaxFilter to attenuate external interferences. Signal space separation (MaxFilter parameters: spatiotemporal signal space separation [SSS], down-sample from 1000Hz to 250Hz, correlation limit between inner and outer subspaces used to reject overlapping intersecting inner/outer signals during spatiotemporal SSS: 0.98, movement compensation using cHPI coils [default step size: 10 ms]) was applied. The data was then converted into Statistical Parametric Mapping (SPM) format and additional pre-processing and analyses were conducted in MATLAB (MathWorks, Natick, MA, USA) using a combination of in-house-built codes (LBPD, https://github.com/leonardob92/LBPD-1.0.git) and the Oxford Centre for Human Brain Activity (OHBA) Software Library (OSL) (https://ohba-analysis.github.io/osl-docs/)(M. Woolrich et al., 2011) which is a freely available software that builds upon Fieldtrip (Oostenveld et al., 2011), FSL (M. W. Woolrich et al., 2009), and SPM (Penny et al., 2007) toolboxes.

The MEG data was visually inspected to identify and remove potential large artifacts. The removed data constituted less than 0.1% of the collected data. For discarding the interference of eyeblinks and heart-beat artefacts from the brain data, independent component analysis (ICA) was used (OSL implementation). The first step was decomposing the original signal into independent components. Then, all the components were correlated with the activity that the ECG and EOG channels recorded. The highly correlated components were also visually inspected, to further validate the accuracy of this procedure. More specifically, their topographic distribution across MEG channels were compared to the typical distribution associated with eyeblink and heartbeat activity to check whether they matched. The components in which the correlations and visual inspections converged were discarded. The third step was rebuilding the signal from the remaining components. Then, the data was low pass filtered (cut-off of 30 Hz) to remove some additional noise and increase the clarity of the brain signal. Finally, the data was epoched (one epoch for each musical sequence). As conceivable, since different sequences had different durations, the length of the epochs was defined with a pre-stimulus time of 100 ms and a post-stimulus time which depended on the overall duration of the musical sequence for each trial (Dataset 1: 125 ms for each tone L epoch length of 3 sec; 350 ms L 3 sec; 650 ms L 4 sec; Dataset 2: 350 ms L 3 sec; 1000 ms L 6 sec; 2000 ms L 11 sec; 5000 ms L 26 sec). Both datasets were then baseline-corrected by subtracting the mean brain signal recorded in the baseline from the brain signal over the entire duration of the epoch.

### Source reconstruction

MEG is an excellent tool for detecting whole-brain activity with high temporal precision. For obtaining a complete picture of the underlying whole-brain activity however, the brain’s spatial component must be accounted for and identified. In this case, the beamforming method was employed. A combination of in-house-built codes and codes available in OSL, SPM, and FieldTrip served as its foundation.

The reconstruction of the brain sources the MEG signal originates from poses an inverse problem, given that the MEG recording shows cortical neural activity, but does not shed light on the brain regions the signal is stemming from. To solve this problem, beamforming algorithms were used, which implemented the following steps: (i) designing a forward model and (ii) computation of the inverse solution.

This theoretical forward model is designed so that it considers each brain source as an active dipole (brain voxel). Its output is describing how the unitary strength of each dipole would be reflected over all MEG sensors. Only the magnetometer channels were employed here, and an 8-mm grid, that ended up returning 3559 dipole locations (voxels) over the entire brain. This was due to differences between magnetometers and gradiometers. The former are the MEG sensors capturing activity originating in deep sources of the brain better, while the latter are stronger in detecting activity stemming from the portion of the cortex in closest proximity to each gradiometer. Since we hypothesised that regions in the medial temporal lobe (e.g. hippocampus) were particularly relevant for long-term auditory recognition, we opted for the magnetometers to compute the source reconstruction. After the individual structural T1 data was co-registered with the fiducial points (i. e., information about head landmarks), the forward model was computed with the adaptation of the widely used method “Single Shell”, that is presented in detail in Nolte, (2003). The output of this computation is referred to as “leadfield model” and was stored in the matrix L (sources x MEG channels). In three cases structural T1 was not available. In these instances, the leadfield computation was performed using a template (MNI152-T1 with 8-mm spatial resolution).

After, the inverse solution was computed. For this - as mentioned above – beamforming was chosen, which is a popular and effective algorithm widely used in the field. This procedure sequentially applies a different set of weights to the source locations, in order to identify the contribution of each source to the activity picked up by the MEG channels. This procedure was done for every time-point of the recorded brain data. The following main steps summarise the beamforming inverse solution.

The following equation (1) describes the data recorded by MEG sensors (*B*) at time *t*:

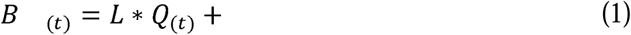

where *L* stands for the leadfield model, Q represents the dipole matrix carrying the activity of each active dipole (q) over time and L is accounting for noise (see Huang and colleagues (1999) for details). For solving the inverse problem, Q must be computed. The first step in the beamforming algorithm is the computation of weights. Then they are applied to the MEG sensors at each time-point. This is shown for the single dipole q in equation (2):

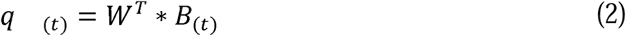

For obtaining *q*, the computation of the weights *W* is necessary (the subscript *T* refers to transpose matrix). To achieve this, the beamforming relies on the matrix multiplication between *L* and the covariance matrix between MEG sensors *C*, which is computed on the concatenated experimental trials. Specifically, for each brain source *q*, the weights *W_q_* are computed as shown in equation (3):

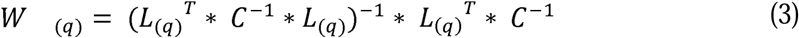

Notably, the leadfield model was computed for the three main orientations of each brain source (dipole), according to Nolte (2003). Prior to the computation of weights, the singular value decomposition algorithm was used on the matrix multiplication reported in equation (4). This was done to reduce the three orientations to one. This method is widely adopted for the simplification for the beamforming output.

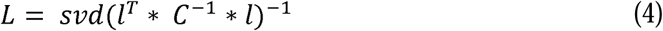

In this case, *L* is the resolved one-orientation model from (3), while *l* stands for the leadfield model with the three orientations. The final step was applying the weights to each brain source and time-point. Notably, the covariance matrix *C* has been computed on the continuous signal, the estimation of which is the result of concatenating the trials of all experimental conditions. The weights were applied to the brain data belonging to each of the conditions and normalised based on Luckhoo and colleagues (2014) to counterbalance the reconstruction bias towards the centre of the head. Then, the weights were applied to the neural activity averaged over trials, independently for each experimental condition. The procedure resulted in a time series for each of the 3559 brain sources for each condition, commonly known as neural activity index. For each brain source, the sign ambiguity of the evoked response time series was adjusted using its sign correspondence with the N100 response to the first tone of the auditory sequences (Bonetti et al., 2024).

### Statistical analysis – neural sources

As described above, the source reconstruction was conducted in an 8-mm space, resulting in 3559 brain voxels, each with a corresponding time series. Here, we were specifically interested in the brain regions of interest (ROIs) primarily recruited during musical long-term recognition. Thus, as shown in **Figure S3**, we selected the six large functional ROIs described by Bonetti et al. (2024) used the same paradigm in a different context. Here, the time series of the brain voxels within each of the six ROIs were averaged, giving rise to only one time series for each ROI. Then, we computed one two-sided t-test contrasting M versus N conditions, independently for each ROI, time-point and speed. To correct for multiple comparisons, we used one-dimensional (1D) cluster-based Monte-Carlo simulations (MCS, α = .05, MCS *p-value* = .001). This procedure, described in detail in Bonetti et al., (2024) consists of binarising the time series based on the significant time-points emerged from the t-tests described above. Here, first the clusters of the contiguous significant time-points are identified. Second, 1000 permutations of the binarised time series are performed. Third, the maximum cluster size obtained for each permutation is extracted and a reference distribution of the cluster sizes for the randomised data is built. Fourth, the clusters in the original data that were larger than 99.9% of the permuted ones were considered significant. This procedure was followed for all the analyses reported in this study which involved neural data. **Figures 3, 4** and **5** and **Table 1** report the results, while **Table S1** show the statistics in detail.

## Data availability

The pre-processed neuroimaging data will be made available upon reasonable request.

## Code availability

The MEG data was first pre-processed using MaxFilter 2.2.15. Then, the data was further pre-processed using Matlab R2016a or later (MathWorks, Natick, Massachusetts, United States of America). Specifically, we used codes from the Oxford Centre for Human Brain Activity Software Library (OSL), FMRIB Software Library (FSL) 6.0, SPM12 and Fieldtrip.

In-house-built code and functions used in this study are part of the LBPD repository which is available at the following link: https://doi.org/10.5281/zenodo.10701724.

The full analysis pipeline used in this study is available at the following link: https://github.com/leonardob92/ConceptGeneralisation_AuditorySequenceRecognition_Speed.git

## Acknowledgements

The Center for Music in the Brain (MIB) is funded by the Danish National Research Foundation (project number DNRF117). L.B. is supported by Lundbeck Foundation (Talent Prize 2022), Carlsberg Foundation (CF20-0239), Center for Music in the Brain, Linacre College of the University of Oxford, and Nordic Mensa Fund. M.L.K. is supported by Center for Music in the Brain and Centre for Eudaimonia and Human Flourishing, which is funded by the Pettit and Carlsberg Foundations. Additionally, we thank Fundación Mutua Madrileña (Mutua Madrileña Foundation, Madrid, Spain) for the support provided to G.F.R. and Mensa: The International High IQ Society (Italian section) for the support provided to F.C.

## Author contributions statement

L.B., G.F.R. and E.B. conceived the hypotheses. L.B. and G.F.R. designed the study. L.B., M.L.K., E.B. and P.V. recruited the resources for the experiment. L.B., G.F.R. and F.C. collected the data. L.B., G.F.R. and S.A.S. performed pre-processing and statistical analysis. E.B., M.L.K. and P.V. provided essential help to interpret and frame the results within the neuroscientific literature. L.B. and S.A.S. wrote the first draft of the manuscript. L.B., G.F.R. and S.A.S. prepared the figures. All the authors contributed to and approved the final version of the manuscript.

## Supplementary material

### Supplementary figures

**Figure S1.**
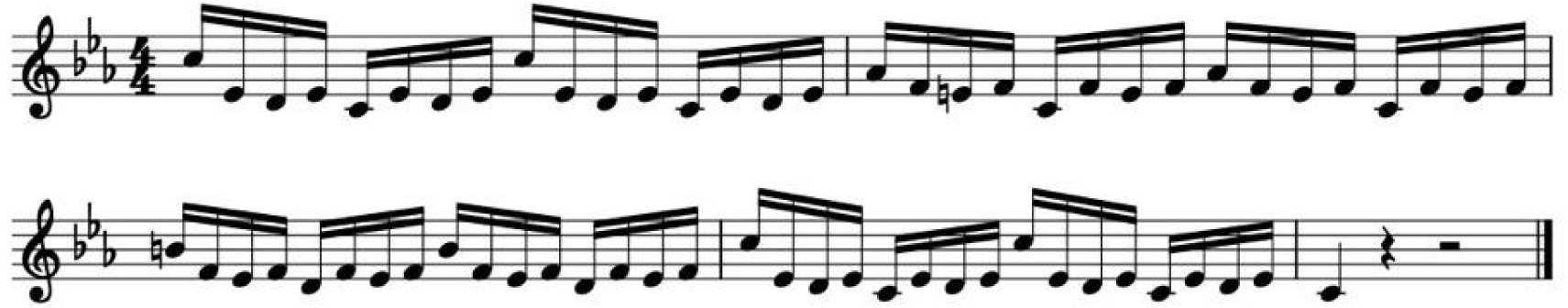
Musical piece used in the encoding phase of the auditory old/new paradigm. In the first part of the auditory old/new paradigm, first participants listened to a short musical piece twice and were asked to memorise it as much as possible. The musical piece consisted of the first four bars of the right-hand part of Johann Sebastian Bach’s Prelude No. 2 in C Minor, BWV 847. In this piece, each bar comprised 16 tones. Thus, the total number of tones was 16*4 = 64. Each tone lasted approximately 350 ms for a total duration of 22400 ms. In addition, to provide a sense of musical closure, we included a final tone after the four bars which lasted 1000 ms. Thus, the total was 23400 ms which correspond to 23.4 seconds. This figure illustrates the piece in musical notation.

**Figure S2.**
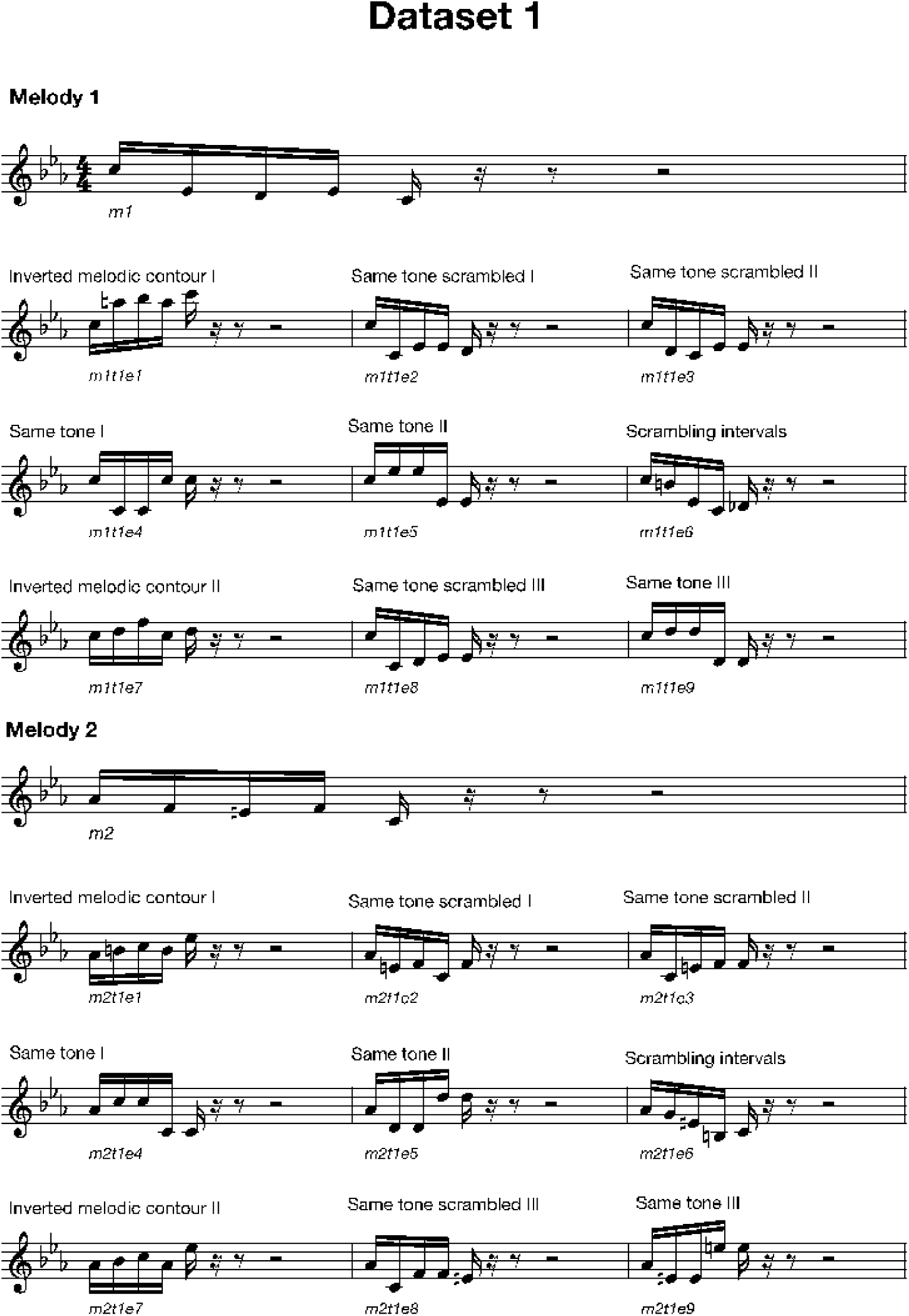
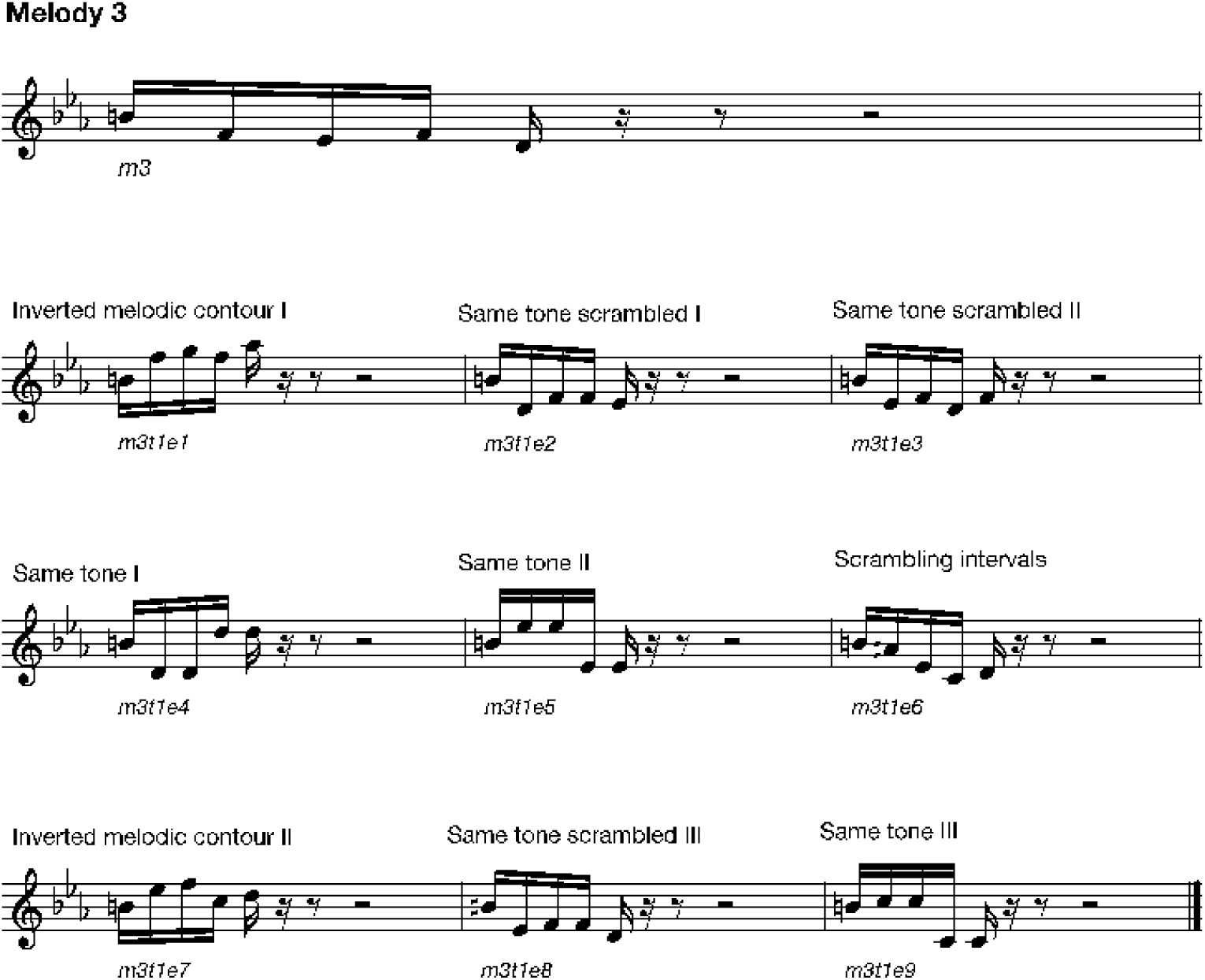
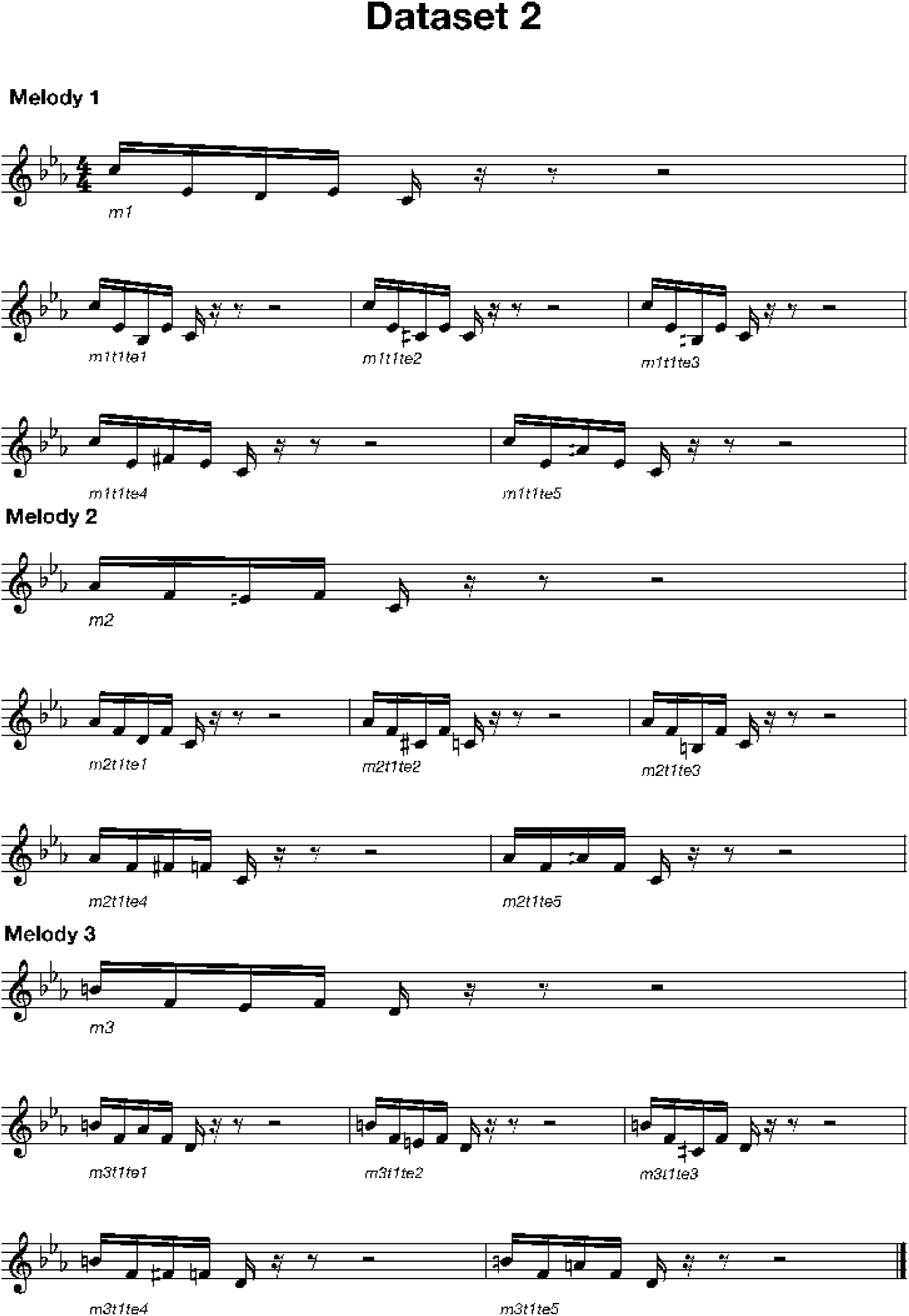
Auditory sequences used in the experiment. The figure shows all auditory sequences used in the experiment in musical notation. For both Datasets 1 and 2, the memorised (M) sequences were three and comprised the first five tones of the first three measures of the musical piece. In Dataset, these three sequences (m1, m2, and m3) were presented nine times each, for a total of 27 trials for the first recognition block (speed 350 ms), and seven times each, for a total of 21 trials, for the second recognition block (speeds 125 ms and 650 ms). In Dataset 2, each melody was repeated four times, for a total of 12 trials for each of the four speeds (350 ms, 1000 ms, 2000 ms and 5000 ms). In Dataset 1, the novel (N) sequences were created through systematic variations of the three M sequences, changing every musical tone of the sequence after the first (NT1) tone. In the first recognition block (speed 250 ms), we created nine variations for each of the original M sequence, resulting in 27 N sequences. As shown in this figure, the nine variations were built according to the following criteria: (i) inverted melodic contours (used twice): the melodic contour of the variation was inverted with regards to the original M sequence (i.e., if the M sequence had the following melodic contour: up-up-down-up, the N sequence would be: down-down-up-down); (ii) same tone scrambled (used three times): the remaining tones of the M sequence were randomly scrambled (e.g., M sequence: C-B-D-B-C, was converted into NT1 sequence: C-C-B-B-D); (iii) same tone (used three times): the same tone was used several times, in some cases changing only the octave (e.g., M sequence: C-B-D-B-C, was transformed into NT1 sequence: C-B-B-B^8^-B^8^); (iv) scrambling intervals (used once): the intervals between the tones were randomly scrambled (e.g., M sequence: 6thM – 2ndM – 2ndm – 5g, was adapted to NT1 sequence: 2ndm, 6thM, 5g, 2ndM). In the second recognition block, seven variations for each of the three M original melodies were randomly selected from the pool of nine variations, resulting in 21 N sequences for each speed (125 ms and 650 ms). In Dataset 2, the N sequences were constructed in a slightly different way compared to Dataset 1. Here, only the tone number three of the sequences was changed and thus the N sequences were named NT3T. Five N sequences were created for each speed (350 ms, 1000 ms, 2000 ms and 5000 ms) and each of the three M sequences, for a total of 60 N sequences.

**Figure S3.**
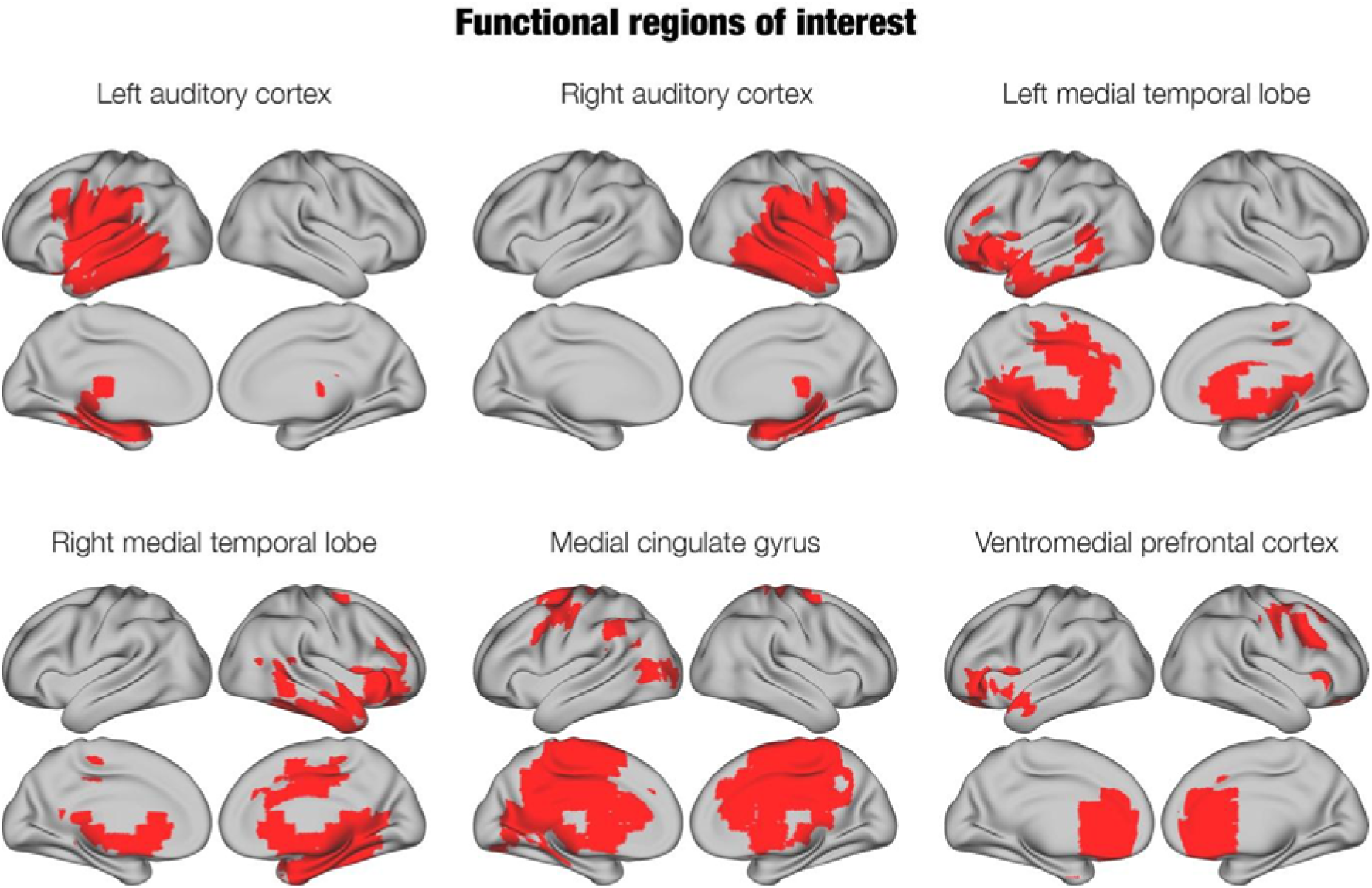
Functional regions of interest (ROIs) derived from the brain activity underlying the task. The main activity during recognition of memorised (M) and novel (N) auditory sequences gave rise to the following six functional ROIs: left (ACL, i) and right auditory cortex (ACR, ii); left (LMTL, iii) and right (RMTL, iv) medial temporal lobe; medial cingulate gyrus (MC, v), and ventromedial prefrontal cortex (VMPFC, vi).

### Supplementary Table

The table is available at the following link: https://docs.google.com/spreadsheets/d/13T1ywM_ntx9kiHNPj0Gsyvol8GD4y_FP/edit?usp=sharing&ouid=102433637013211014958&rtpof=true&sd=true

***Table S1. Source-localised differences in evoked responses across experimental conditions and speeds***

*Univariate analysis computed independently on the six Regions of Interest (ROIs) of the functional parcellation: left auditory cortex (ACL), right auditory cortex (ACR), left medial temporal lobe and inferior temporal cortex (LMTL), right medial temporal lobe and inferior temporal cortex (RMTL), medial cingulate (MC), ventromedial prefrontal cortex (VMPFC). Here, for each ROI, two-sided t-tests were conducted independently for each time-point, experimental conditions (memorised [M] versus novel T1 [NT1] for Dataset 1 and memorised [M] versus novel T3T [NT3T] for Dataset 2) and speed and corrected for multiple comparisons using one-dimensional (1D) cluster-based Monte-Carlo simulations (MCS;* α *= .05, MCS p-value = .001). The significant clusters outputted by the Monte-Carlo simulations (MCS) are reported showing cluster size, p-value, temporal extent of the cluster and peak t-value within the cluster*.

## References

Bertelson, P., & Tisseyre, F. (1970). Perceiving the sequence of speech and non-speech stimuli. Quarterly Journal of Experimental Psychology, 22(4), 653–662. 10.1080/14640747008401943

Bonetti, L., Brattico, E., Bruzzone, S. E. P., Donati, G., Deco, G., Pantazis, D., Vuust, P., & Kringelbach, M. L. (2023). Brain recognition of previously learned versus novel temporal sequences: A differential simultaneous processing. Cerebral Cortex, 33(9), 5524–5537.

Bonetti, L., Brattico, E., Carlomagno, F., Donati, G., Cabral, J., Haumann, N. T., Deco, G., Vuust, P., & Kringelbach, M. L. (2021). Rapid encoding of musical tones discovered in whole-brain connectivity. NeuroImage, 245, 118735.

Bonetti, L., Fernández-Rubio, G., Carlomagno, F., Dietz, M., Pantazis, D., Vuust, P., & Kringelbach, M. L. (2024). Spatiotemporal brain hierarchies of auditory memory recognition and predictive coding. Nature Communications, 15(1), 4313. 10.1038/s41467-024-48302-4

Bonetti, L., Brattico, E., Carlomagno, F., Cabral, J., Stevner, A., Deco, G., Whybrow, P. C., Pearce, M., Pantazis, D., & Vuust, P. (2024). Spatiotemporal brain dynamics during recognition of the music of Johann Sebastian Bach. Cerebral Cortex, 34 (8), bhae320. 10.1093/cercor/bhae320

Bowman, C. R., & Zeithamova, D. (2018). Abstract Memory Representations in the Ventromedial Prefrontal Cortex and Hippocampus Support Concept Generalization. Journal of Neuroscience, 38(10), 2605–2614. 10.1523/JNEUROSCI.2811-17.2018

Brattico, E., & Delussi, M. (2024). Making sense of music: Insights from neurophysiology and connectivity analyses in naturalistic listening conditions. Hearing Research, 441, 108923. 10.1016/j.heares.2023.108923

Bronstein, D. (2016). Aristotle on Knowledge and Learning: The Posterior Analytics. Oxford University Press. 10.1093/acprof:oso/9780198724902.001.0001

Criscuolo, A., Pando-Naude, V., Bonetti, L., Vuust, P., & Brattico, E. (2022). An ALE meta-analytic review of musical expertise. Scientific Reports, 12(1), 11726. 10.1038/s41598-022-14959-4

Croonen, W. L. M. (1994). Effects of length, tonal structure, and contour in the recognition of tone series. Perception & Psychophysics, 55(6), 623–632. 10.3758/BF03211677

Dowling, W. J., Bartlett, J. C., Halpern, A. R., & Andrews, M. W. (2008). Melody recognition at fast and slow tempos: Effects of age, experience, and familiarity. Perception & Psychophysics, 70(3), 496–502. 10.3758/PP.70.3.496

Dunsmoor, J. E., & Murphy, G. L. (2015). Categories, concepts, and conditioning: How humans generalize fear. Trends in Cognitive Sciences, 19(2), 73–77. 10.1016/j.tics.2014.12.003

Fernández-Rubio, G., Brattico, E., Kotz, S. A., Kringelbach, M. L., Vuust, P., & Bonetti, L. (2022). Magnetoencephalography recordings reveal the spatiotemporal dynamics of recognition memory for complex versus simple auditory sequences. Communications Biology, 5(1), 1272.

Fernández-Rubio, G., Carlomagno, F., Vuust, P., Kringelbach, M. L., & Bonetti, L. (2022). Associations between abstract working memory abilities and brain activity underlying long-term recognition of auditory sequences. PNAS Nexus, 1(4), pgac216.

Fernández-Rubio, G., Olsen, E. R., Klarlund, M., Mallon, O., Carlomagno, F., Vuust, P., Kringelbach, M. L., Brattico, E., & Bonetti, L. (2024). Investigating the impact of age on auditory short-term, long-term, and working memory. Psychology of Music, 52(2), 187–198. 10.1177/03057356231183404

Freedman, D. J., & Assad, J. A. (2016). Neuronal Mechanisms of Visual Categorization: An Abstract View on Decision Making. Annual Review of Neuroscience, 39(1), 129–147. 10.1146/annurev-neuro-071714-033919

Frixione, M., & Lieto, A. (2012). Prototypes Vs Exemplars in Concept Representation: Proceedings of the International Conference on Knowledge Engineering and Ontology Development, 226–232. 10.5220/0004139102260232

Geirhos, R., Temme, C. R. M., Rauber, J., Schütt, H. H., Bethge, M., & Wichmann, F. A. (2020). Generalisation in humans and deep neural networks (arXiv:1808.08750). arXiv. http://arxiv.org/abs/1808.08750

Halpern, A. R., Zatorre, R. J., Bouffard, M., & Johnson, J. A. (2004). Behavioral and neural correlates of perceived and imagined musical timbre. Neuropsychologia, 42(9), 1281–1292. 10.1016/j.neuropsychologia.2003.12.017

Hantschel, F., & Bullerjahn, C. (2016). The Use of Prototype Theory for Understanding the Perception and Concept Formation of Musical Styles (pp. 151–156).

Hasson, U., Chen, J., & Honey, C. J. (2015). Hierarchical process memory: Memory as an integral component of information processing. Trends in Cognitive Sciences, 19(6), 304–313. 10.1016/j.tics.2015.04.006

Huang, M. X., Mosher, J. C., & Leahy, R. M. (1999). A sensor-weighted overlapping-sphere head model and exhaustive head model comparison for MEG. Physics in Medicine & Biology, 44(2), 423. 10.1088/0031-9155/44/2/010

Höhne, J., & Tangermann, M. (2012). How stimulation speed affects Event-Related Potentials and BCI performance | IEEE Conference Publication | IEEE Xplore. https://ieeexplore.ieee.org/abstract/document/6346300?casa_token=5rmfteDP9ygAAAAA:lTYfOU3_SblYj8fVLtfPZSnXdPYA0RG1guwdGsWGedNxMKvOuc5IwLtEpbj6jP5E4ewEXql7IQ

Kaernbach, C. (2004). The Memory of Noise. Experimental Psychology, 51(4), 240–248. 10.1027/1618-3169.51.4.240

Kiefer, M., & Pulvermüller, F. (2012). Conceptual representations in mind and brain: Theoretical developments, current evidence and future directions. Cortex, 48(7), 805–825. 10.1016/j.cortex.2011.04.006

Koelsch, S. (2018). Investigating the Neural Encoding of Emotion with Music. Neuron, 98(6), 1075–1079. 10.1016/j.neuron.2018.04.029

Koelsch, S. (2020). A coordinate-based meta-analysis of music-evoked emotions. NeuroImage, 223, 117350. 10.1016/j.neuroimage.2020.117350

Ladefoged, P., & Broadbent, D. E. (1960). Perception of Sequence in Auditory Events. Quarterly Journal of Experimental Psychology, 12(3), 162–170. 10.1080/17470216008416720

Lerner, Y., Honey, C. J., Katkov, M., & Hasson, U. (2014). Temporal scaling of neural responses to compressed and dilated natural speech. Journal of Neurophysiology, 111(12), 2433–2444. 10.1152/jn.00497.2013

Luckhoo, H. T., Brookes, M. J., & Woolrich, M. W. (2014). Multi-session statistics on beamformed MEG data. NeuroImage, 95, 330–335. 10.1016/j.neuroimage.2013.12.026

McKeown, D., Mills, R., & Mercer, T. (2011). Comparisons of Complex Sounds across Extended Retention Intervals Survives Reading Aloud. Perception, 40(10), 1193–1205. 10.1068/p6988

Millidge, B., Seth, A., & Buckley, C. L. (2022). Predictive Coding: A Theoretical and Experimental Review (arXiv:2107.12979). arXiv. 10.48550/arXiv.2107.12979

Morey, R. A., Haswell, C. C., Stjepanović, D., Dunsmoor, J. E., & LaBar, K. S. (2020). Neural correlates of conceptual-level fear generalization in posttraumatic stress disorder. Neuropsychopharmacology, 45(8), 1380–1389. 10.1038/s41386-020-0661-8

Nolte, G. (2003). The magnetic lead field theorem in the quasi-static approximation and its use for magnetoencephalography forward calculation in realistic volume conductors. Physics in Medicine & Biology, 48(22), 3637. 10.1088/0031-9155/48/22/002

Oostenveld, R., Fries, P., Maris, E., & Schoffelen, J.-M. (2011). FieldTrip: Open source software for advanced analysis of MEG, EEG, and invasive electrophysiological data. Computational Intelligence and Neuroscience, 2011, 156869. 10.1155/2011/156869

Pando-Naude, V., Patyczek, A., Bonetti, L., & Vuust, P. (2021). An ALE meta-analytic review of top-down and bottom-up processing of music in the brain. Scientific Reports, 11(1), 20813.

Pearl, J. (2018). What is Gained from Past Learning. Journal of Causal Inference, 6(1). 10.1515/jci-2018-0005

Penny, W., Friston, K., Ashburner, J., Kiebel, S., & Nichols, T. (2007). Statistical Parametric Mapping: The Analysis of Functional Brain Images. 10.1016/B978-0-12-372560-8.X5000-1

Rosch, E., & Lloyd, B. B. (Eds.). (1978). Cognition and Categorization. Routledge. 10.4324/9781032633275

Schulze, K., & Tillmann, B. (2013). Working memory for pitch, timbre, and words. Memory, 21(3), 377–395. 10.1080/09658211.2012.731070

Schulze, K., Jay Dowling, W., & Tillmann, B. (2012). Working Memory for Tonal and Atonal Sequences during a Forward and a Backward Recognition Task. Music Perception, 29(3), 255–267. 10.1525/mp.2012.29.3.255

Sharaev, M., Andreev, A., Artemov, A., Burnaev, E., Kondratyeva, E., Sushchinskaya, S., Samotaeva, I., Gaskin, V., & Bernstein, A. (2018). Pattern Recognition Pipeline for Neuroimaging Data. In L. Pancioni, F. Schwenker, & E. Trentin (Eds.), Artificial Neural Networks in Pattern Recognition (pp. 306–319). Springer International Publishing. 10.1007/978-3-319-99978-4_24

Sommer, T. (2016). The Emergence of Knowledge and How it Supports the Memory for Novel Related Information. Cerebral Cortex, bhw031. 10.1093/cercor/bhw031

Surprenant, A. M., Brown, M. A., Jalbert, A., Neath, I., Bireta, T. J., & Tehan, G. (2011). Backward Recall and the Word Length Effect. The American Journal of Psychology, 124(1), 75–86. 10.5406/amerjpsyc.124.1.0075

Sutojo, S., Thiemann, J., Kohlrausch, A., & van de Par, S. (2020). Auditory Gestalt Rules and Their Application. In J. Blauert & J. Braasch (Eds.), The Technology of Binaural Understanding (pp. 33–59). Springer International Publishing. 10.1007/978-3-030-00386-9_2

Tehan, G., & Mills, K. (2007). Working memory and short-term memory storage: What does backward recall tell us. Working Memory: Behavioural and Neural Correlates, 153–163.

Terhardt, E. (2016). Gestalt Principles and Music Perception. In Auditory Processing of Complex Sounds (pp. 157–166). Routledge. 10.4324/9781315622347-19

Tillmann, B. (2012). Music and Language Perception: Expectations, Structural Integration, and Cognitive Sequencing. Topics in Cognitive Science, 4(4), 568–584. 10.1111/j.1756-8765.2012.01209.x

Vaidya, A. R., Jones, H. M., Castillo, J., & Badre, D. (2021). Neural representation of abstract task structure during generalization. eLife, 10, e63226. 10.7554/eLife.63226

Van de Cruys, S., & Wagemans, J. (2011). Putting Reward in Art: A Tentative Prediction Error Account of Visual Art. I-Perception, 2(9), 1035–1062. 10.1068/i0466aap

Warren, R. M., Gardner, D. A., Brubaker, B. S., & Bashford, J. A. (1991). Melodic and Nonmelodic Sequences of Tones: Effects of Duration on Perception. Music Perception, 8(3), 277–290.

Williamson, V. J., Baddeley, A. D., & Hitch, G. J. (2010). Musicians’ and nonmusicians’ short-term memory for verbal and musical sequences: Comparing phonological similarity and pitch proximity. Memory & Cognition, 38(2), 163–175. 10.3758/MC.38.2.163

Woolrich, M., Hunt, L., Groves, A., & Barnes, G. (2011). MEG beamforming using Bayesian PCA for adaptive data covariance matrix regularization. NeuroImage, 57(4), 1466–1479. 10.1016/j.neuroimage.2011.04.041

Woolrich, M. W., Jbabdi, S., Patenaude, B., Chappell, M., Makni, S., Behrens, T., Beckmann, C., Jenkinson, M., & Smith, S. M. (2009). Bayesian analysis of neuroimaging data in FSL. NeuroImage, 45(1 Suppl), S173-186. 10.1016/j.neuroimage.2008.10.055

Wu, J., Zeng, W., & Yan, F. (2018). Hierarchical Temporal Memory method for time-series-based anomaly detection. Neurocomputing, 273, 535–546. 10.1016/j.neucom.2017.08.026

